# Activated interstitial macrophages are a predominant target of viral takeover and focus of inflammation in COVID-19 initiation in human lung

**DOI:** 10.1101/2022.05.10.491266

**Authors:** Timothy Ting-Hsuan Wu, Kyle J. Travaglini, Arjun Rustagi, Duo Xu, Yue Zhang, Leonid Andronov, SoRi Jang, Astrid Gillich, Roozbeh Dehghannasiri, Giovanny Martínez-Colón, Aimee Beck, Daniel Dan Liu, Aaron J. Wilk, Maurizio Morri, Winston L. Trope, Rob Bierman, Irving L. Weissman, Joseph B. Shrager, Stephen R. Quake, Christin S. Kuo, Julia Salzman, W. E. Moerner, Peter S. Kim, Catherine A. Blish, Mark A. Krasnow

## Abstract

Early stages of deadly respiratory diseases such as COVID-19 have been challenging to elucidate due to lack of an experimental system that recapitulates the cellular and structural complexity of the human lung while allowing precise control over disease initiation and systematic interrogation of molecular events at cellular resolution. Here we show healthy human lung slices cultured *ex vivo* can be productively infected with SARS-CoV-2, and the cellular tropism of the virus and its distinct and dynamic effects on host cell gene expression can be determined by single cell RNA sequencing and reconstruction of “infection pseudotime” for individual lung cell types. This revealed that the prominent SARS-CoV-2 target is a population of activated interstitial macrophages (IMs), which as infection proceeds accumulate thousands of viral RNA molecules per cell, comprising up to 60% of the cellular transcriptome and including canonical and novel subgenomic RNAs. During viral takeover of IMs, there is cell-autonomous induction of a pro-fibrotic program (*TGFB1*, *SPP1*), and an inflammatory program characterized by the early interferon response, chemokines (*CCL2*, 7, *8*, *13, CXCL10*) and cytokines (*IL6, IL10)*, along with destruction of cellular architecture and formation of dense viral genomic RNA bodies revealed by super-resolution microscopy. In contrast, alveolar macrophages (AMs) showed neither viral takeover nor induction of a substantial inflammatory response, although both purified AMs and IMs supported production of infectious virions. Spike-dependent viral entry into AMs was neutralized by blockade of ACE2 or Sialoadhesin/CD169, whereas IM entry was neutralized only by DC-SIGN/CD209 blockade. These results provide a molecular characterization of the initiation of COVID-19 in human lung tissue, identify activated IMs as a prominent site of viral takeover and focus of inflammation and fibrosis, and suggest therapeutic targeting of the DC-SIGN/CD209 entry mechanism to prevent IM infection, destruction and early pathology in COVID-19 pneumonia. Our approach can be generalized to define the initiation program and evaluate therapeutics for any human lung infection at cellular resolution.

## INTRODUCTION

Lower respiratory infections are one of the leading causes of death worldwide^1,2^, accelerated by the current Coronavirus Disease 2019 (COVID-19) pandemic^3^. Most such infections, including COVID-19, start innocuously in the upper respiratory tract and become dangerous when they reach alveoli^4-9^, the site of gas exchange, but the critical transition to life threatening pneumonia and acute respiratory distress syndrome (ARDS) has been difficult to elucidate. For practical and ethical reasons, such early and key steps in human pathogenesis have been inferred, with rare exception^10,11^, from examination of late or end stage patient lung lavage, biopsy, or autopsy specimens, using classical histopathological methods^12-14^ and recently single cell multi-omic profiling^15-20^.

The above approaches provide a picture of COVID-19 pneumonia at unprecedented cellular and molecular resolution, and have suggested models of pathogenesis involving not only infection of the alveolar epithelium but also implicating alveolar capillaries, macrophages and other myeloid cells^18,19,21-24^ and production of various inflammatory cytokines and chemokines^15,19^. It remains unclear which cells are the direct virus targets in the human lung, and the nature of their virus-induced host response – in particular, the origin and sequence of molecular signals that initiate, sustain, and propagate the inflammatory cascade that leads to COVID-19 ARDS^4^.

These early pathogenic events hold the key to understanding and preventing the transitions to the deadly and systemic forms of COVID-19, but we know little about them. This is due to difficulty accessing human lung tissue at this critical transition, and the sheer number of lung (>58) cell types potentially involved. This cellular complexity has made pathogenic mechanisms challenging to empirically address even in the most sophisticated human lung organoid systems^25-28^ and animal models^5-7,20,29-33^.

Here we describe an experimental model of SARS-CoV-2 infection that allows systematic interrogation of the early molecular events and pathogenic mechanism of COVID-19 at cellular resolution in native human lung tissue. We determine the cellular tropism of SARS-CoV-2 and its distinct and dynamic effects on host cell gene expression for individual lung cell types. The most prominent targets are two lung resident macrophage populations, one of which the virus takes over the transcriptome and induces a specific host interferon anti-viral program along with seven chemokines, and pro-inflammatory, as well as pro-fibrotic cytokines that can signal to a diverse array of lung immune and structural cell types. We propose that this early focus of lung inflammation is an important step in the transition to the deadly and systemic forms of COVID-19 and a potential new therapeutic target.

## RESULTS

### Human lung slices cultured *ex vivo* are productively infected by SARS-CoV-2

To define the early events of SARS-CoV-2 infection in human lung, we cut thick sections (∼300*–*500 µm “slices”) of fresh lung tissue procured from therapeutic surgical resections or organ donors, and placed the slices in culture medium containing DMEM/F12 and 10% FBS (Fig. 1a). We then exposed them to SARS-CoV-2 (USA-WA1/2020) at a multiplicity of infection (MOI) of 1 for two hours, then allowing infection to proceed for 24 or 72 hours. Plaque assay of culture supernatants demonstrated production of infectious virions that increased between 24 and 72 hours of culturing (Fig. 1b, Extended Data Fig.1). Productive infection was abrogated by pre-inactivation of the viral stocks with heat or ultraviolet (UV)-C, or by treatment of the cultures with 10 µM remdesivir, an RNA-dependent RNA polymerase inhibitor used as a COVID-19 therapeutic (Fig. 1b).

**Figure 1.**
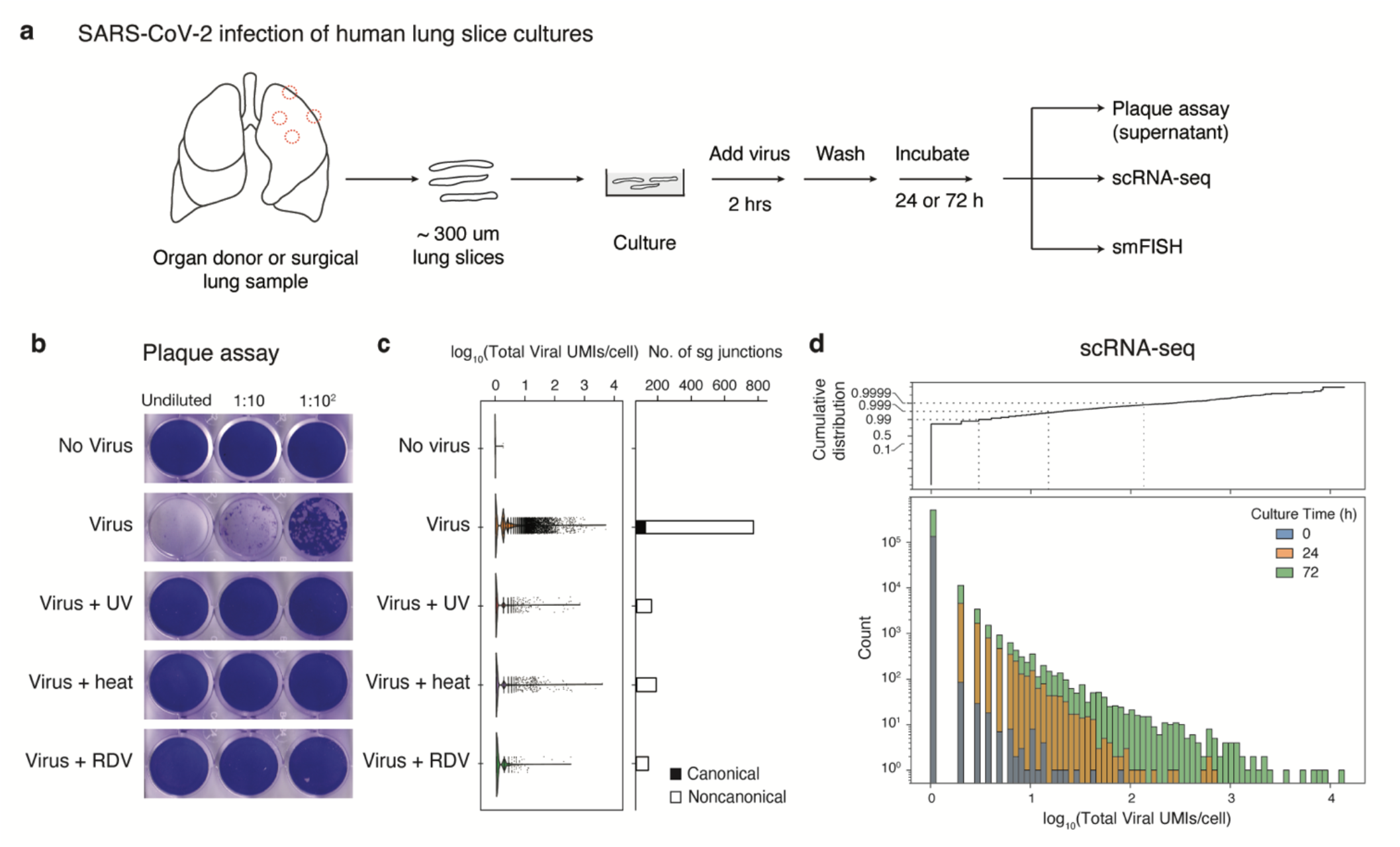
Detection of virion production, viral RNA amplification, and subgenomic RNA in cultured human lung tissue infected ex vivo by SARS-CoV-2. (a) Strategy for slicing, culturing, infecting, and analyzing human lung tissue from healthy, surgically resected or organ donor lungs. In each case, distal and proximal lung regions were sampled and sliced into 300-500 um sections. Slices were cultured (DMEM/F12 medium supplemented with 10% FBS) at 37°C and subsequently exposed to SARS-CoV-2 for 2 hours, washed to remove free virus, and cultured in supplemented DMEM/F12 for 24 or 72 hours to allow infection to proceed before assaying supernatant for virion production by plaque assay, preserving tissue in 10% NBF for histological staining and multiplex single molecule fluorescence hybridization (smFISH), or dissociating tissue for 10x single cell RNA sequencing (scRNA-seq). (b) Productive infection of lung slices from case 5 measured by plaque assay. Lung slices were mock-infected for 72 hours (“No virus”) or infected with purified SARS-CoV-2/WA1 virions (estimated multiplicity of infection ∼1, see Methods) without pretreatment of the virus (“Virus”) or controls with virus pretreated with ultraviolet-C light (“Virus + UV”) or heat (“Virus + heat”) to inactivate virus, or with the virus-infected culture treated with the viral RdRp inhibitor remdesivir at 10μM final concentration (“Virus + RDV”). The supernatant was then harvested and plaque assay performed on VeroE6 cells. (c) scRNA-seq analysis of cultured lung slices from case 5 infected with virus and indicated control conditions as in (b). Violin plot (left) shows viral RNA expression levels (Total number of Unique Molecular Identifiers (UMIs) for detected viral RNAs) in single cells, and bar plot (right) shows number of viral subgenomic RNA junctions detected by SICILIAN^43^. Canonical, transcription-regulatory sequence (TRS) mediated junctions from the 5’ leader (TRS-L) to the 5’ end of open reading frames in the gene body (TRS-B); noncanonical, all other subgenomic junctions detected that pass SICILIAN statistical test. (d) Bar graph (bottom) showing dynamic range of viral RNA molecules expressed (Total number of viral UMIs/cell) in profiled single cells (Count) from scRNA-seq of infected lung slice cultures from all cases as in (a) but from lung slices cultured as indicated for 0, 24, or 72 hours following exposure to SARS-CoV-2. Dashed lines (in Cumulative distribution, top), expression levels for 99%, 99.9% and 99.99% of profiled cells.

To characterize viral and host gene expression during SARS-CoV-2 infection, slices were dissociated and analyzed by single-cell RNA sequencing (scRNA-seq, 10x Genomics), adapting the methods we and others previously used to construct a comprehensive transcriptomic atlas of the healthy human lung^34-40^ to capture, quantify, and map SARS-CoV-2 viral gene expression along with host gene expression in each profiled lung cell^41,42^. The number of viral RNA molecules detected per infected cell spanned a wide range (Fig.1c, d), with the vast majority (∼99%) of profiled cells from infected lung slices containing few or no detected viral RNA molecules (Fig. 1d). But the rest of the cells (∼1%) expressed tens to hundreds of viral RNA molecules per cell at 24 hours, and by 72 hours the distribution had shifted to even higher values with rare cells (∼0.01%) accumulating thousands of viral RNAs per cell (Fig. 1d), paralleling the increase in virus production during this period (Extended Data Fig.1). As with infectious virions, viral RNA levels determined by scRNA-seq were diminished by heat or UV-C inactivation of the virus stocks, or by treatment of the cultures with remdesivir (Fig. 1c).

We also investigated the junctional structure and processing of the viral RNA molecules by analyzing our scRNA-seq dataset using the SICILIAN framework^43^, which identifies RNA sequencing reads that map discontinuously in a genome, such as reads that span splice junctions of eukaryotic mRNAs or the subgenomic junctions of the nested SARS-CoV-2 mRNAs. We detected canonical subgenomic junctions among the rare sequence reads outside their 3’ ends, confirming generation of canonical SARS-CoV-2 mRNAs in the lung slice cultures (Fig. 1c, right panel). In addition, we identified dozens of novel subgenomic junctions, indicating widespread generation of diverse non-canonical subgenomic viral RNAs along with canonical subgenomic forms during lung infection (Fig. 1c, Extended Data Fig. 2, Table S1). These non-canonical junctions included three that spliced the standard viral 5’ leader sequence to a novel downstream site, as well as 494 junctions between two novel internal sites in the genome and 479 junctions between an internal and 3’ site (the most abundant non-canonical species detected, consistent with the strong 3’ end bias of 10x 3.1 technology). Some of these non-canonical RNAs are predicted to encode novel viral proteins, or alter potential regulatory sequences in the 3’ non-coding region of the viral mRNA. Heat or UV-C inactivation, or remdesivir treatment each abrogated formation of both canonical and non-canonical junctions (Fig. 1c, right panel). Together, these data demonstrate that lung cultures support ongoing, productive viral gene expression and replication.

**Figure 2.**
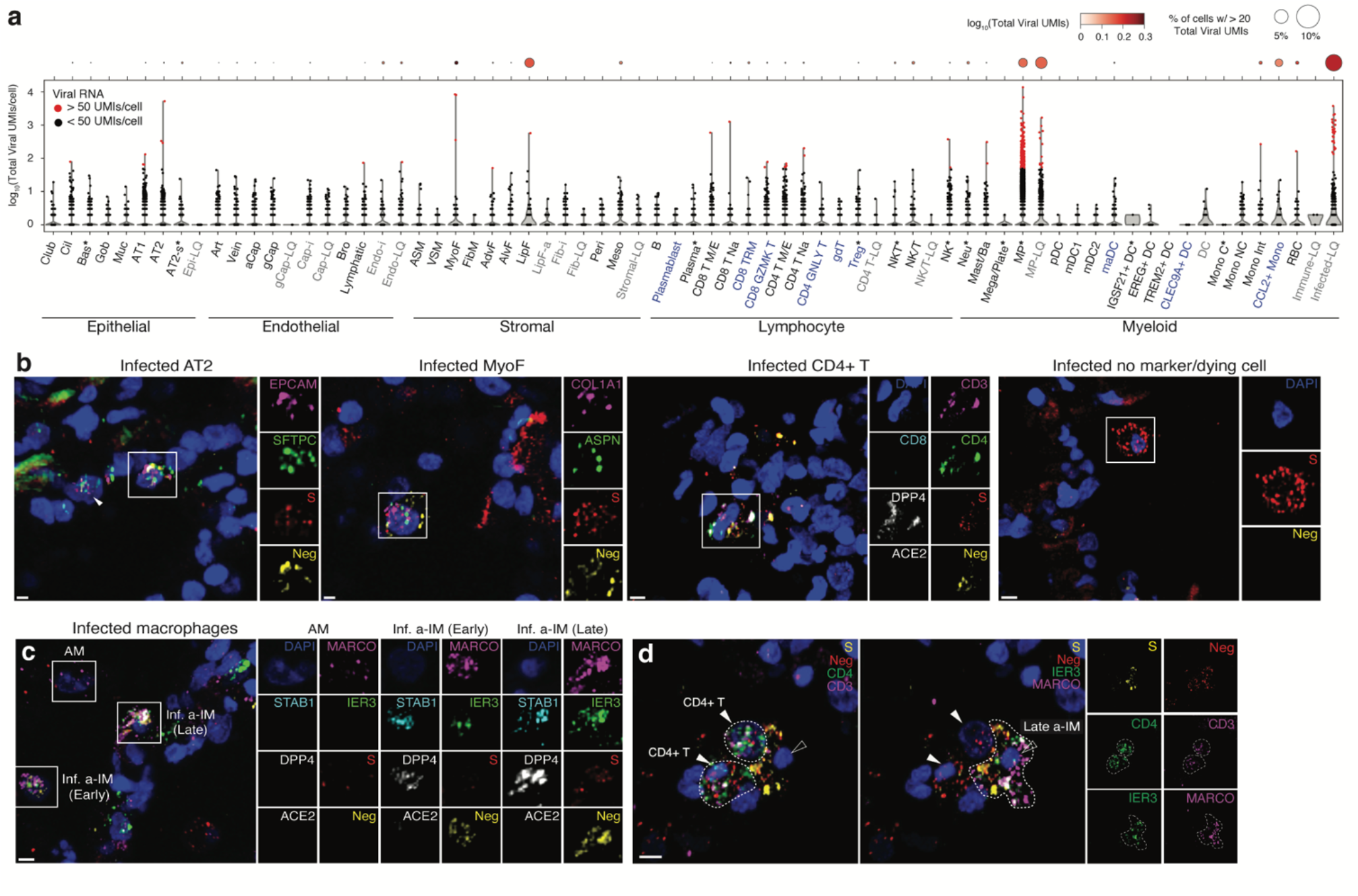
A comprehensive map of SARS-CoV-2 cell tropism in the human lung. (a) Violin plot of viral RNA expression level (log10-transformed viral UMIs) in the single cells of each of the molecular cell types detected by scRNA-seq of the lung slice infections from cases 1-4. Dot plot above shows pseudo-bulk viral RNA expression level for each cell type; dot size indicates the percentage of cells in each type with detected expression of viral RNA (thresholded at ≥20 viral UMI), and shading shows mean level of expression for the cells that passed detected expression threshold. Asterisk, cell types in which a proliferative subpopulation was detected but merged with the non-proliferating population in this plot (note these include basal, macrophage, and NKT cells, none of which were previously found to include a proliferating subpopulation in the native lung); blue text, additional cell types not detected or annotated in our native human lung cell atlas^34^; grey text, cell types only observed in cultured lung slices. (b) RNAscope multiplex single molecule fluorescence in situ hybridization (smFISH) of infected lung slice culture from case 2, fixed 72 hours after infection, showing close-ups (boxed, split channels at right) of canonical (alveolar epithelial type 2 (AT2)) and novel (myofibroblast (MyoF), CD4 T cell) lung cell targets of the virus, as well as an infected cell at a late stage of infection as indicated by high expression of positive strand viral RNA detected with S probe (red) and little or no expression of negative strand viral RNA (Neg, yellow) or the cell type markers examined. Probes were: positive strand viral RNA (viral S, red), negative strand viral RNA (Neg, antisense viral orf1ab, yellow), the canonical SARS-CoV-2 receptor *ACE2* (white), compartment markers for the epithelium (*EPCAM*, magenta), stroma (*COL1A1*, magenta), and cell type markers identifying alveolar epithelial type 2 (AT2) cells (*SFTPC*, green), myofibroblasts (MyoF; *ASPN*, green), CD4 T cells (*CD3*, magenta; *CD4*, green; *CD8*, cyan). (c) RNAscope smFISH of lung slice cultures as above detecting infected macrophage subtypes: viral S (red), negative strand RNA (antisense Orf1ab, yellow), the canonical SARS-CoV-2 receptor *ACE2* (white) and a receptor (*DPP4*) used by the related MERS coronavirus (white), general macrophage marker *MARCO* (magenta), and activated interstitial macrophage (a-IM) markers *STAB1* (cyan) and *IER3* (green). Close-ups of boxed regions (right) show alveolar macrophages (AMs, *MARCO*^+^*STAB1*^-^*IER3*^-^) that express few S puncta and no negative puncta, and activated interstitial macrophages (a-IMs, *MARCO*^+^*STAB1*^-^*IER3^+^*) in early infection (“early a-IM”) expressing few S puncta and abundant negative puncta, and a-IMs in late infection (“late a-IM”) with abundant S and negative puncta. (d) RNAscope smFISH detecting interaction between infected a-IM (*MARCO*^+^*IER3*^+^) expressing viral S and negative strand RNA (antisense Orf1ab), and two CD4 T cells (*CD3*^+^*CD4*^+^) expressing viral negative strand RNA but not viral S. Split panels at right show individual channels. Scale bars, 10 µm.

### A cellular atlas of SARS-CoV-2 tropism in the human lung

The cellular tropism of a virus — the set of host cells that allow viral entry and replication — is among the most characteristic and significant determinants of virulence. Historically tropism has been inferred from autopsy specimens, often weeks, months, or even years after disease onset. More recently, tropism has been predicted from expression patterns of entry receptors identified by biochemical or functional screening in heterologous cell types^44^. For SARS-CoV-2, a small subset of lung epithelial types (AT2, ciliated, AT1, club, and goblet cells) were predicted to be the major direct targets for SARS-CoV-2 based on their expression of the canonical SARS-CoV-2 receptor ACE2 and protease TMPRSS2^18,25,34,45^. However, studies of COVID-19 autopsy lungs have detected viral gene products in various epithelial and endothelial cells, fibroblasts, and myeloid cells, indicating widespread viral presence at least in end-stage disease^18,19^.

To determine SARS-CoV-2 lung cell tropism empirically and directly compare infection of lung cell types in their natural context, we first used the most sensitive and specific markers from our molecular atlas of the healthy human lung^34^ to identify the cell types present in the cultured lung slices from their transcriptomic profiles, then assessed their viral RNA levels in the infected cultures. Of the 176,382 cells with high quality transcriptomes obtained from infected lung slices of four donor lungs, along with those of the 112,359 cells from mock-infected slices (cultured without viral addition) and 95,389 uncultured control cells (directly from freshly-cut lung slices), we identified 55 distinct molecular lung cell types distributed across the major tissue compartments (Fig. 2a, Extended Data Fig. 3, Table S2). These included most (46 out of 58, 80%) of the cell types described in the healthy human lung^34^ plus 5 additional types of lymphocytes (e.g., CD4+ cytotoxic T lymphocytes, γδ T cells, regulatory T cells, tissue-resident memory CD8+ T cells, GZMK+CD8+ T cell; Fig. 2a, blue) along with culture-induced proliferative states of signaling alveolar type 2 (AT2-s) cells, NK cells, and dendritic cells (DCs) and several culture-induced proliferative and activation states of fibroblasts, which could not be ascribed to any previously defined fibroblast types (Fig. 2a, grey). The only cell types not recovered after culturing were rare myeloid types (e.g., IGSF21+ DCs, TREM2+ DCs, classical monocytes), which may egress from the slices or not survive during culture (Extended Data Fig.3, Table S2).

**Figure 3.**
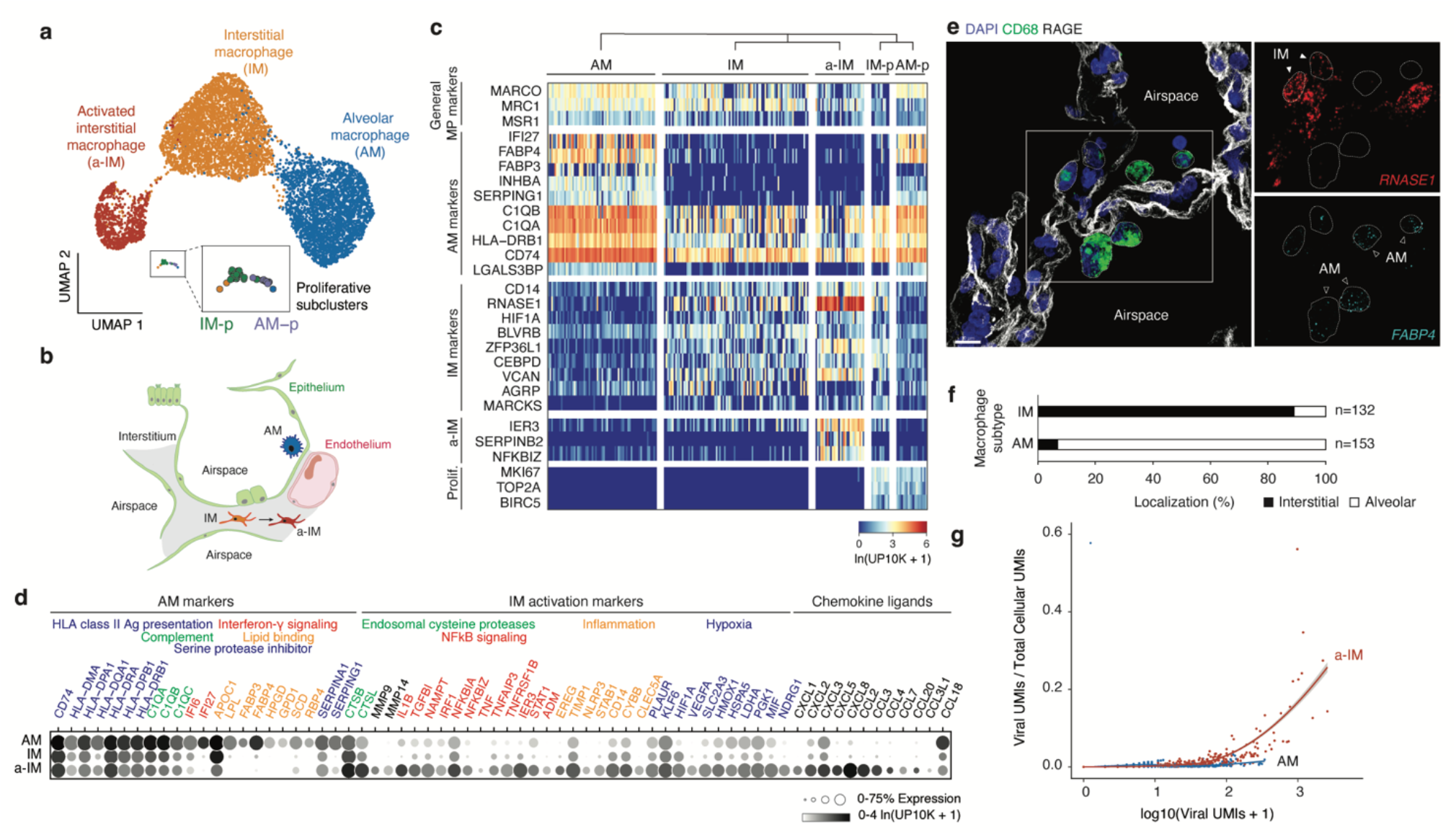
Identity, tissue localization, and viral takeover of molecularly distinct macrophage populations in the human lung. (a) Uniform Manifold Approximation and Projection (UMAP) projection of molecularly distinct macrophage subpopulations in cultured human lung slices from cases 1 and 4 identified by computational clustering of their individual 10x scRNA-seq expression profiles (colored dots). Note three major molecular types: alveolar macrophages (AM) and newly designated (see panel e) interstitial macrophages (IM) and activated interstitial macrophages (a-IM), plus a minor cluster of proliferating macrophages (boxed) that using distinguishing markers shown in panel c could be subclassified as proliferative AMs (AM-p) or proliferative IMs (IM-p) (expanded box). (b) Schematic of alveoli, with epithelial barrier (green) comprised of AT1, AT2, and AT2-s cells, and endothelial barrier of underlying capillary comprised of aerocytes and general capillary cells. AMs reside in the airspace, while IMs and a-IMs reside in the interstitium (grey) bounded by the basal surfaces of epithelium and endothelium of neighboring alveoli. (c) Heatmap of expression of general macrophage marker genes (rows) in the individual macrophages from (a) (columns) of the indicated subtypes (for visualization, randomly downsampled to < 80 cells), and top differentially expressed genes that distinguish the subtypes. Note all clusters express general macrophage marker genes, but each has its own set of selectively expressed markers. (d) Dot plot showing fraction of expressing cells and mean expression (among expressing cells) of AM markers and IM activation markers in the macrophage subtypes from (a). Encoded proteins with related functions are indicated by color of the gene names. (e) Tissue localization of macrophage subtypes by RNAscope single molecule fluorescence in situ hybridization (smFISH) and immunostaining in control, non-cultured human lung from case 2. Markers shown: general macrophage antigen CD68 (green, protein), AT1 antigen RAGE (white, protein), AM marker FABP4 (cyan, RNA), and IM marker RNASE1 (red, RNA). Scale bar, 30 µm. Note AMs localized to the apical side of AT1 cells that comprise alveolar epithelium (interpreted to be alveolar airspace), whereas IMs localized to the basal side of AT1 cells and are bounded by epithelium (interpreted to be the interstitial space). (f) Quantification of anatomical localization of AMs and IMs. Cells with substantial (>80%) colocalization with RAGE AT1 antigen were scored as interstitial, and those without substantial colocalization with RAGE AT1 antigen (<20% to account for AMs contacting the apical side of AT1 cells, as schematized in b) and any other cells were scored as alveolar. (g) Viral RNA takeover of the host transcriptome (Viral UMIs/ Total Cellular UMIs) graphed against viral expression (Total Viral UMIs) in single cells of AMs (blue dots) and a-IMs (red dots) from the infected human lung slices from case 1. Note that beginning at ∼70 viral RNA molecules (UMIs) per cell, viral RNA begins to rapidly increase to thousands of viral molecules per cell and dominate (“takeover”) the host cell transcriptome (25-60% total cellular UMIs) in a-IMs, whereas in AMs viral RNA never exceeded a few hundred UMI per cell and 1-2% of the host transcriptome, even at corresponding viral RNA cellular loads.

Cellular SARS-CoV-2 viral RNA levels across the 55 human lung cell types in the infected cultures are shown in Fig. 2a. Although 10-20 viral RNA molecules were detected in about one-third of the molecular cell types in the infected cultures, cells with high viral levels (hundreds to thousands of SARS-CoV-2 UMI per cell) were rare and restricted to six cell types. One was AT2 cells, the primary predicted lung target of SARS-CoV-2^34,46^. The others included myofibroblasts, lipofibroblasts, two molecular types of T cells and NK cells, and macrophages. Macrophages were the most prominent lung targets, accounting for 75% of cells with 50 or more viral UMI per cell. However, even for macrophages, such cells represented only a small proportion of the recovered cell type (0.5% of all macrophages), indicating inefficient entry or a sensitive subpopulation (see below). One caveat to this tropism analysis is that identities could not be assigned to 16% of cells with 50 or more viral UMI per cell because they did not robustly express cell type markers (“unidentified” cell types, Fig. 2a), presumably due to downregulation of the host transcriptome and cell destruction during viral takeover. Most cells with high viral load were detected in cultures at 72 but not 24 hours after infection, indicating that the intervening 48 hours is the critical period of viral RNA amplification in most lung cell types.

To validate these lung cell tropism results, visualize the infected cells, and localize foci of viral replication, we performed multiplexed single molecule fluorescence *in situ* hybridization (smFISH) of the infected lung slices to simultaneously detect positive strand viral RNA (S gene probe), negative strand viral RNA (replication intermediate, Orf1ab gene probe), the canonical viral receptor ACE2, and markers of the infected cell types detected in scRNA-seq (Fig. 2b,c). We found both positive and negative strand viral RNA in AT2 cells (*SFTPC*^+^*EPCAM*^+^), myofibroblasts (*ASPN*^+^*COL1A2*^+^), macrophages (*PTPRC*^+^*MARCO*^+^) and exceedingly rarely, CD4 T cells (*PTPRC*^+^*CD3*^+^*CD4*^+^). We also detected cells filled with viral mRNA molecules but no negative strand RNA (the early replication intermediate), or any of the cell-type markers in our panel; these are likely cells in the terminal, lytic stage of infection. Infected cells were generally scattered throughout the infected lung tissue, but rare clusters were detected such as an infected macrophage associated with two CD4 T cells (Fig. 2d).

For AT2 cells, myofibroblasts, and T cells, the cells with high viral load were rare in the tissue sections, as in the scRNA-seq tropism analysis. In contrast, infected macrophages were more abundant and showed a broad and seemingly continuous range of viral RNA molecules. Some macrophages (*PTPRC*^+^*MARCO*^+^) showed a few (1-3 puncta) positive-strand viral RNA molecules but no negative strand viral RNA, whereas others expressed a few (1-3 puncta) negative strand viral RNA molecules alongside a wide range (1 to dozens of puncta) of positive strand viral RNA molecules (Fig. 2c).

### SARS-CoV-2 takeover of an activated interstitial macrophage subtype

We reasoned that the macrophages in the lung slice cultures with SARS-CoV-2 RNA levels spanning several orders of magnitude — from tens to thousands of UMIs in the scRNA-seq analysis (Fig. 2a) and from one to dozens of puncta detected by smFISH (Fig. 2c) — were infected cells harboring active intermediates that had progressed to different stages of infection, those with highest RNA loads having progressed furthest in the infection cycle. This is consistent with our finding that cells harvested 72 hours after infection generally had higher viral RNA levels than those harvested at 24 hours (Fig. 1d).

To resolve the apparent heterogeneity in the infected macrophages, we further clustered the gene expression profiles of macrophages in the lung slices and found that they separated into three distinct clusters (Fig. 3a). One had higher expression of genes involved in functions ascribed to mature alveolar macrophages, including antigen-presentation major histocompatibility complex class II (MHCII) genes (*HLA-DPA1*, *HLA-DPB1*, *HLA-DQA1*, *HLA-DQB1*, and *CD74*) and genes involved in lipid homeostasis (*LPL*, *APOC1*, *FABP3*, *FABP4*, and *HPGD*; Fig. 3c,d)^47^. smFISH showed cells expressing these markers were larger and rounder in morphology than the others, and localized to the alveolar airspace (Fig. 3b,e,f), hence we refer to them as “alveolar macrophages” (AMs). Another cluster we call “interstitial macrophages” (IMs) expressed lower levels of the classical AM markers including *LPL*, *APOC1*, *FABP3*, *FABP4*, and *HPGD* but were enriched for a different set of genes including the monocyte marker *CD14* (Fig. 3c,d), and localized interstitially (Fig. 3b,e,f, Extended Data Fig. 4). The third cluster was transcriptionally similar to interstitial macrophages but also expressed genes known to be activated by NF-κB signaling (*NFKBIA*, *NFKBIZ*, and *IL1B*), inflammation (*IER3*, *EREG*, *TIMP1*, *STAB1*), and hypoxia-induced factor *HIF1A*; we call them “activated interstitial macrophages” (Fig. 3c,d, a-IMs). Although a-IMs were detected in the uncultured control lung slices, they were a minor population. However, upon culturing, almost all IMs became activated; similar populations of activated lung macrophages have been observed infiltrating tumors in the intact human lung^48,49^ and in other inflammatory conditions^50,51^.

**Figure 4.**
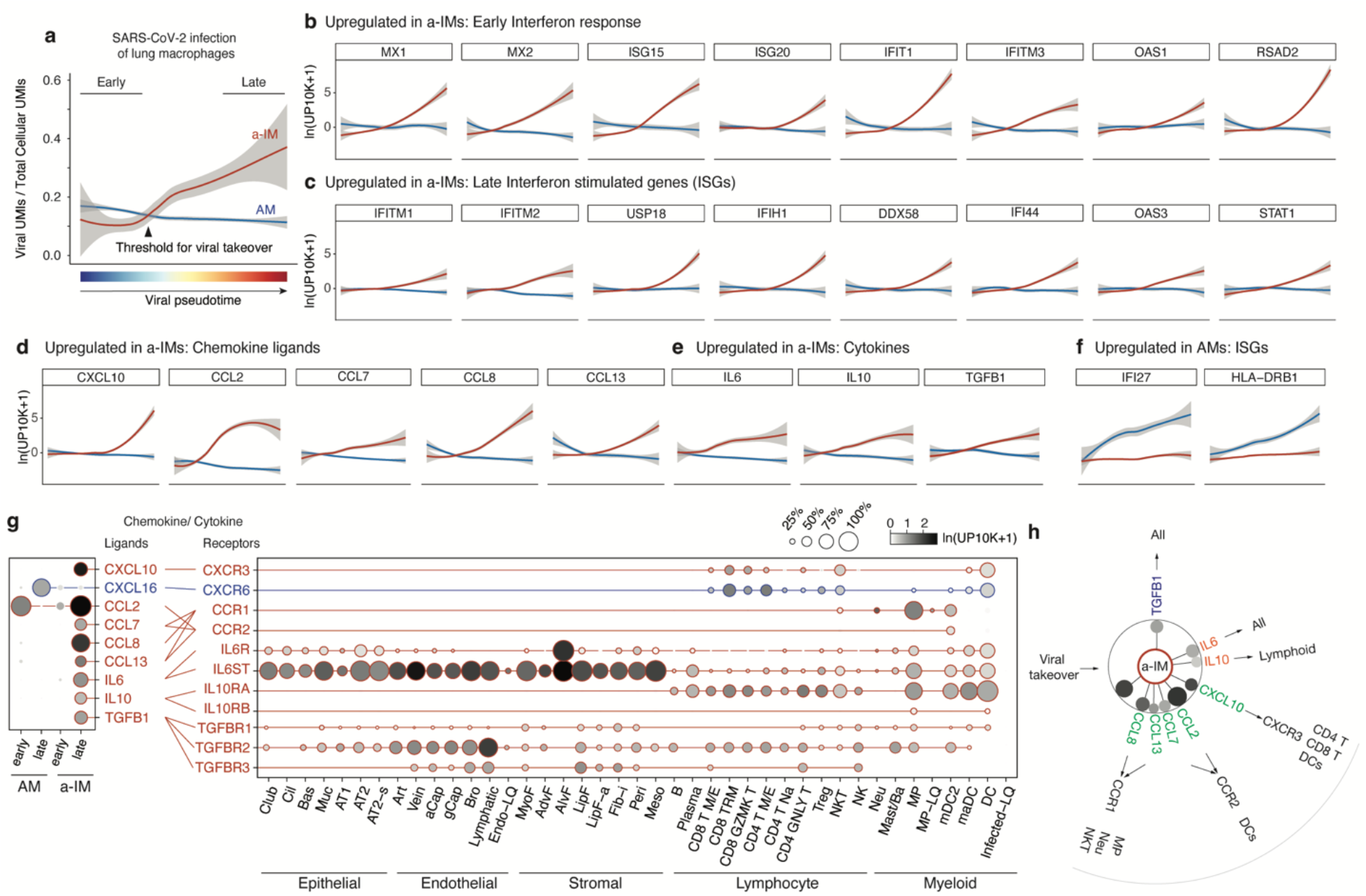
Differential induction of host response and inflammatory genes in activated interstitial and alveolar macrophages shown by infection pseudotime. (a) Viral takeover (Viral UMIs/ Total Cellular UMIs) graphed against viral infection pseudotime for AMs (blue) and a-IMs (red) from the infected human lung slices from case 1; grey shading indicates 95% confidence interval. Pseudotime was separately computed for AMs and a-IMs by taking a linear combination of principal components that best correlated with monotonic increase in viral expression, then linearly rescaling between 0-1. “Early” cells in each infection pseudotime trajectory were defined by normalized pseudotime < 0.2, and “Late” cells were defined by normalized pseudotime > 0.8. (b-f) Host gene expression profiles of AMs and a-IMs plotted along infection pseudotime, as in (a). A differential expression test was performed on the top 250 genes with the highest loadings for the infection pseudotime axis, and the selected genes presented (b-f) (visualized for the AMs and a-IMs in infected human lung slices from case 1) were among those that had statistically significant association with infection pseudotime as indicated. (b) Early interferon response genes that were significantly associated with a-IM pseudotime trajectory. (c) Late interferon stimulated genes (ISGs) that were significantly associated with a-IM pseudotime trajectory. (d) Chemokine ligands that were significantly associated with a-IM pseudotime trajectory. (e) Cytokines that were significantly associated with a-IM pseudotime trajectory. (f) ISGs that were significantly associated with AM pseudotime trajectory. (g) Dot plots (left) showing discretized expression of chemokine/cytokine ligands that were differentially expressed between early and late pseudotime a-IMs (*CXCL10*, *CCL2*, *CCL7*, *CCL8*, *CCL13*, *IL6*, *IL10, TGFB1*) and AMs (*CXCL16*), and their cognate receptors (right) in human lung cells (all infected conditions from cases 1-4); only cell types and chemokines with detected expression are shown. Lines connect ligands with cognate receptor. Red, virally induced in a-IMs; blue, differentially virally induced in AMs. (h) Summary schematic depicting the six cytokine and chemokine genes induced in a-IMs during viral takeover (dot sizes scaled to percentage expression and shaded with mean expression as in g), and the lung cell targets of the encoded inflammatory signals predicted from expression of the cognate receptor genes. Outbound arrows from a-IMs, cytokine signaling to lung cell targets or chemokine recruitment of immune cells toward a-IMs.

Comparison of SARS-CoV-2 infection of the macrophage subtypes in the slice cultures revealed a striking difference in viral RNA accumulation in AMs vs. a-IMs. Although both AMs and a-IMs could accumulate hundreds of viral RNAs, only in a-IMs did viral RNA accumulate beyond 300 viral UMI per cell and result in viral domination (“takeover”) of the host cell transcriptome (Fig. 3g). Viral takeover reached up to 60% of an a-IM transcriptome (ratio of viral to total UMIs in a cell), whereas it never exceeded 2% of an AM transcriptome (Fig. 3g). Thus, in a-IMs, SARS-CoV-2 can infect and amplify its RNA until it dominates the host transcriptome, whereas viral RNA takeover does not occur in AMs.

### Infection pseudotime of activated interstitial macrophages reveals an early focus of lung inflammation and fibrosis

To characterize the host cell response during viral takeover, we computationally ordered the infected macrophages according to the principal components that best correlated with viral RNA levels and takeover to reconstruct what we refer to as “infection pseudotime” (Fig. 4a, Extended Data Fig. 5), similar to developmental pseudotime^52,53^, providing a dynamic view of the viral gene expression program and its effect on the host transcriptome.

**Figure 5.**
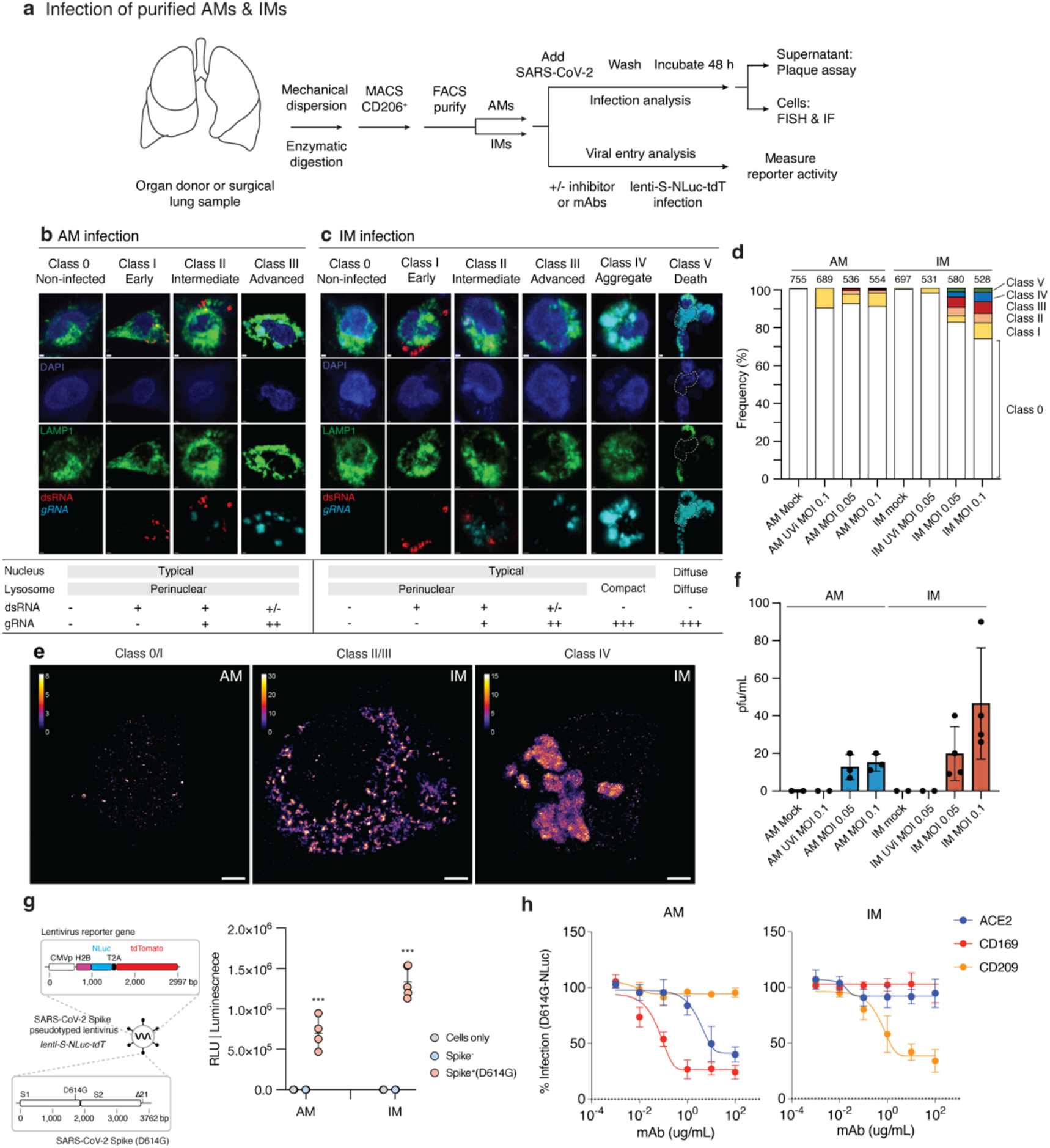
SARS-CoV-2 entry, replication, and productive infection of purified AMs and IMs. (a) Strategy for purification, culture, and infection of human lung macrophages with SARS-CoV-2 virions or a SARS-CoV-2 Spike-pseudotyped lentivirus. Human lung tissue obtained from surgical resections or organ donors were dissociated fresh, then enriched for macrophages by magnetic activated cell separation (MACS) using antibodies against the general lung macrophage antigen CD206, followed by specific AM or IM purification using fluorescent activated cell sorting (FACS) for distinguishing markers (Extended Data Fig. 9). Purified Ams (CD206^+^CD204^hi^) or IMs (CD206^+^CD204^lo^) were cultured at 37°C in DMEM/F12 medium with 10% FBS, and either infected with SARS-CoV-2 virions and analyzed as indicated (panels b-f), or tested for viral entry and the effect of inhibitors and mABs using a SARS-CoV-2 Spike-pseudotyped lentivirus lenti-S-NLuc-tdT that encodes a Nanoluciferase (NLuc) reporter (panels g, h). For SARS-CoV-2 infections, purified AMs or IMs were mock-infected or exposed for 2 hours to untreated or UV-inactivated (UVi) SARS-CoV-2 virions (MOI 0.05 or 0.01), washed to remove free virions, and infection continued for 48 hours before assaying supernatant for virion production by plaque assay or analyzing the infected cells by fluorescence in situ hybridization (FISH using hybridization chain reaction, HCR) and immunofluorescence (IF) staining. (b, c) Infection intermediates and morphologies of SARS-CoV-2 infected AMs (b) or IMs (c) generated as above and then fixed and IF-labeled for lysosomal antigen LAMP1 (green) and the infection dsRNA intermediate (red), followed by HCR for viral genomic RNA (light blue) and DAPI nuclear counterstain (dark blue). Examples of the observed infection classes are shown and their features summarized at panel bottom: Class 0 (non-infected), no expression of either dsRNA or viral gRNA; Class I (early infection): expression of dsRNA only; Class II (intermediate infection), co-expression of dsRNA and viral gRNA; Class III (advanced infection), expression of viral gRNA only; Class IV (aggregates), expression of globular viral gRNA bodies; Class V (cell destruction/death): weak or non-staining of DAPI nuclear stain and LAMP1, and expression of viral RNA. Scale bars, 1 µm. (d) Quantification of (b,c) showing relative abundance of each infection class. Values above each bar, number of cells scored per condition. (e) Super-resolution microscopy of viral gRNA for class 0/1 AM, class II/III IM, and class IV IM from (b, c). Note the large, globular viral gRNA aggregates (“RNA bodies”) throughout the cytoplasm in the class IV IM. Scale bars, 2 µm. (f) SARS-CoV-2 virions released into the medium by the above infected AMs or IMs, as determined by plaque assay of the indicated culture supernatants on a monolayer of VeroE6 cells. pfu, plaque forming units. (g) Viral entry into AMs and IMs depends on SARS-CoV-2 Spike. (Left) Diagram of lenti-S-NLuc-tdT, a lentivirus pseudotyped to express full length SARS-CoV-2 Spike, encoding both S1 and S2 subunits from the D614G variant (Spike+, D614G) protein on its surface and also engineered to express the reporter gene (boxed) encoding nuclear-targeted nanoluciferase (H2B-NLuc) and tdTomato fluorescent protein, separated by a self-cleaving T2A peptide. (Right) lenti-S-NLuc-tdT (Spike+ (D614G)) or a non-pseudotyped control lentivirus (Spike-) were added to purified AMs or IMs in culture, and, after 4 hours, free virions were washed off and infections continued for 48 hours before quantification of infection by expression level (luminescence) of the NLuc reporter. Uninfected AMs or IMs (Cells only) served as background control. RLU, relative light units. NLuc luciferase values are presented as mean ± s.d from two independent experiments, with values normalized to control (non-neutralized) viral infections in each plate. P values are computed from comparing Spike+(D614G) to Spike-controls. ***, P<0.001. (h) Neutralization of lenti-S-NLuc-tdT entry into purified AMs (left) or IMs (right) by the indicated blocking antibodies against ACE2, CD169, or CD209.

Differential gene expression analysis^54^ of a-IMs along infection pseudotime identified host gene expression changes that correlated with viral RNA levels (Table S3); the kinetics of induction of individual genes in infection pseudotime is shown in Fig. 4b-f. A specific set of antiviral genes was upregulated during viral amplification, including the earliest, interferon regulatory factor 3 (IRF3)-dependent type I interferon response genes (*ISG15*, *ISG20*, *IFIT1*, *IFITM3*, *OAS1*, *RSAD2*, *MX1*, *MX2*; Fig. 4b)^55-58^ and many additional canonical interferon-stimulated genes (ISGs; *IFI44*, *IFI44L*, *IFIH1*, *LPAR6*, *USP18*, *HELZ2*, *IFITM1*, *IFITM2*, *STAT1*, *DDX58*, *OAS3*, *XAF1*; Fig. 4c)^59^. This appears to be the cell intrinsic response to infection, presumably resulting from detection of accumulating viral RNA. Viral amplification in a-IMs was also associated with induction of five chemokines (*CCL2*, *CCL7*, *CCL8*, *CCL13*, *CXCL10*; Fig. 4d)) and cytokines *IL10*, and *IL6*, which is among the molecules central to COVID-19 cytokine storm^60^, as well as *TGFB1*, the central mediator of fibrogenesis (Fig. 4e), as well as other genes implicated in pro-fibrotic function (*SPP1, GADD45B*, *ITGB3*, *IGFBP4*; Table S3). In contrast, expression of chemokines *CXCL1*,*2,3,5*, and *8* was downregulated (Table S3).

Alveolar macrophages showed a distinct and more limited response to the virus (Table S3, Fig. 4f). During AM infection pseudotime, only a handful of genes were specifically induced, including *APOC1*, *FDX1*, *IFI27*, *HLA-DRB1*, serine proteases *SERPINA1* and *SERPING1,* and *CXCL16*. Expression of nearly all other chemokines (*CXCL1*, *CXCL2*, *CXCL3*, *CXCL5*, *CXCL8*, *CCL3*, *CCL4*, *CCL20*) was downregulated in AMs.

To predict the cellular targets of the inflammatory and pro-fibrotic signals induced by SARS-CoV-2 infection of a-IMs, we used the single cell gene expression profiles of the infected lung slices to produce a map of cells expressing the cognate receptors (Fig. 4g). Viral induction of *CXCL10* in a-IMs predicts communication to and recruitment of broad classes of CD4 and CD8 T cells via the cognate receptor *CXCR3*, consistent with our observation by smFISH that T cells interacted directly with infected a-IMs. (Fig. 2d). Viral induction of *CCL2* predicts recruitment of specific DC subtypes (mature DCs, mDC2) through *CCR2*, and induction of *CCL8* could recruit neutrophils and create a self-amplifying circuit with macrophages via *CCR1* (Fig. 4h). Viral induction of *TGFB1* predicts pro-fibrotic signaling to most epithelial and fibroblasts expressing both *TGFBR1* and *TGFBR2* (Fig. 4g), including myofibroblasts, which constitute the fibroblast foci in lung fibrosis.

The viral induction of *IL6* and *IL10* along infection pseudotime (Fig. 4g) indicates that infected a-IMs can broadcast potent pro-inflammatory (*IL6*) signals to most other cell types of the lung (broadly expressed, *IL6R* and *IL6ST*/gp130), but the viral induction of the anti-inflammatory cytokine *IL10,* whose receptor (*IL10RA* and *IL10RB*) is mainly expressed in lymphoid cells, and to a lesser extent, myeloid cells, may limit activation of adaptive immunity while enhancing innate inflammation.

Whereas infected AMs restrict viral RNA amplification and generally suppress their communication to other immune cell types, we conclude that infection and takeover induces an early anti-viral cell intrinsic response that is specific to a-IMs (Extended Data Fig. 6) and creates a robust immune and fibrosis signaling center and focus of inflammation (Extended Data Fig. 7) in SARS-CoV-2 infection. Inflammasome activation, recently implicated in severe COVID-19 ^23,24^, was rare and only detected late in a-IM infection (Extended Data Fig. 8, ASC specks).

**Figure 6.**
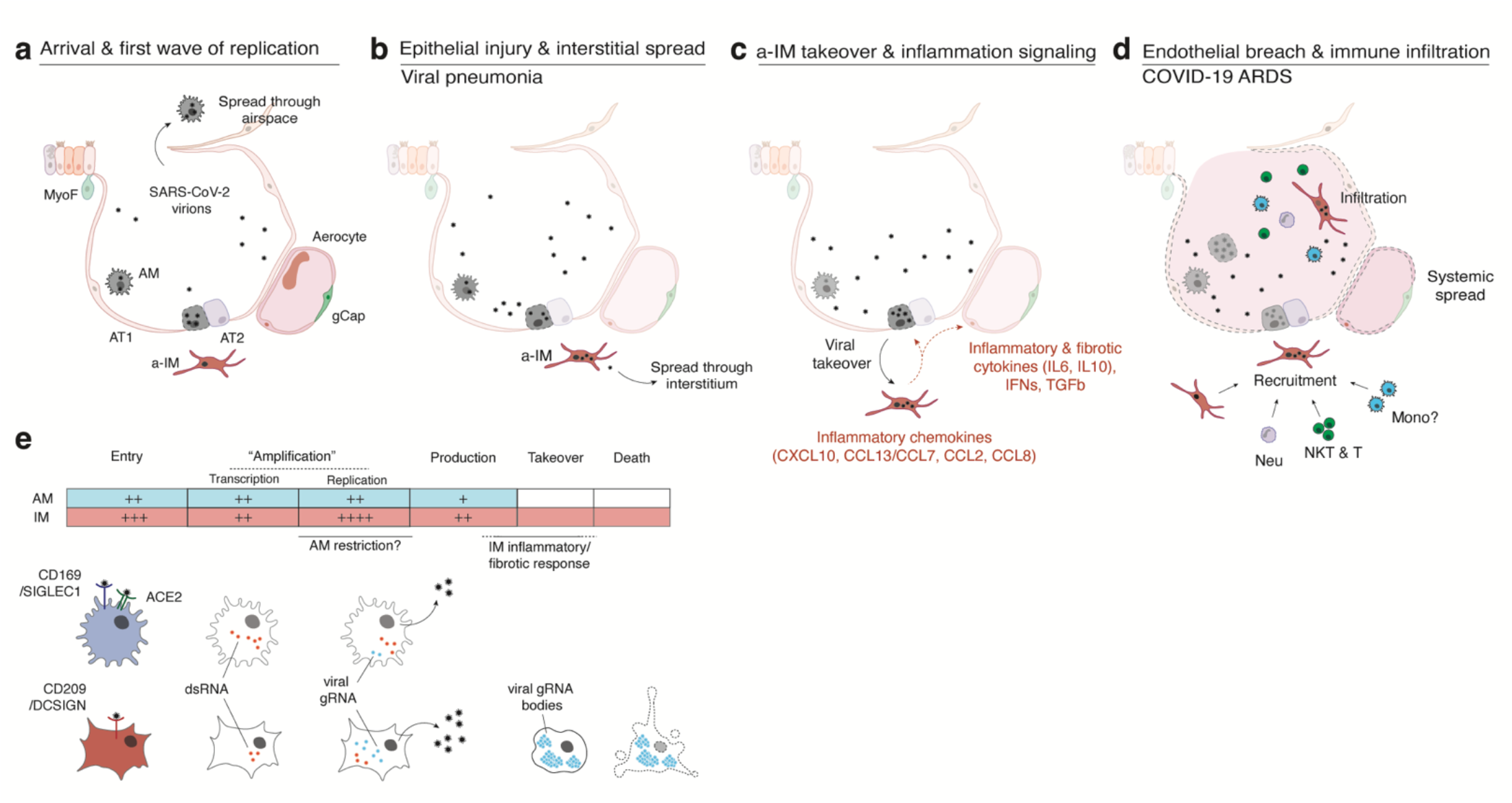
Model of initiation, transition, and pathogenesis of COVID-19 and the viral lifecycle in AMs and IMs. (a-d) Model of COVID-19 initiation in the human lung and transition from viral pneumonia to lethal COVID-19 ARDS. (a) SARS-CoV-2 virion dissemination and arrival in the alveoli. Luminal alveolar macrophages (AM) encounter virions shed from the upper respiratory tract that enter the lung. AMs can express low to moderate numbers of viral RNA molecules and can propagate the infection but “contain” the viral RNA from taking over the total transcriptome and show only a very limited host cell inflammatory response to viral infection. (b) Replication and epithelial injury. SARS-CoV-2 virions enter AT2 cells through ACE2, its canonical receptor, and “replicate” to high viral RNA levels, producing infectious virions and initiating viral pneumonia. (c) a-IM takeover and inflammation signaling. SARS-CoV-2 virions spread to the interstitial space through either transepithelial release of virions by AT2 cells or injury of the epithelial barrier, and enter activated interstitial macrophages (a-IMs). Infected a-IMs can express very high levels of viral RNA that dominate (“take over”) the host transcriptome and can propagate the infection. Viral takeover triggers induction of the chemokines and cytokines shown, forming a focus of inflammatory and fibrotic signaling. (d) Endothelial breach and immune infiltration. The a-IM inflammatory cytokine IL6 targets structural cells of the alveolus causing epithelial and endothelial breakdown, and the inflammatory cytokines recruit the indicated immune cells from the interstitium or bloodstream, which flood and infiltrate the alveolus causing COVID-19 ARDS. Local inflammatory molecules are amplified by circulating immune cells, and reciprocally can spread through the bloodstream to cause systemic symptoms of cytokine storm. (e) Comparison of the SARS-CoV-2 viral lifecycle in AMs and IMs. Although both can produce infectious virions, note differences in viral entry receptors (AMs can use ACE2 and CD169/SIGLEC1, whereas IMs use CD209); viral RNA transcription of dsRNA intermediates (greater in AMs); replication of full-length genomic RNA (greater in IMs); viral takeover, formation of RNA bodies, and induction of a robust host cell inflammatory response (only in IMs), and cell destruction/death (only in IMs).

### SARS-CoV-2 productive infection and destruction of interstitial macrophages

To test whether SARS-CoV-2 can productively infect purified AMs and IMs in isolation and compare the infection cycles, we developed a method for purification of each of these macrophage populations directly from freshly dissociated human lung using sensitive and specific cell surface antigens that distinguish them (Fig. 5a, scheme, Extended Data Fig. 9). The purified AMs (CD206^+^CD204^hi^ cells) and IMs (CD206^+^CD204^lo^ cells) survived in culture for up to a week. We exposed purified AM or IMs to SARS-CoV-2 (USA-WA1/2020) for two hours at a multiplicity of infection (MOI) of 0.05 or 0.1 for two hours, washed to remove unbound virus, then allowed the infections to proceed for 48 hours. Plaque assay of culture supernatants demonstrated production of infectious virions in both populations (Fig. 5f). Although we cannot exclude transient uptake and release of infectious virions into the supernatant, such a mechanism would not explain the observed amplification of viral RNA (see below). Pre-inactivation of viral stocks with UV-C abrogated productive infection of both, confirming a requirement for active replication.

To compare the cellular course of viral replication in AMs or IMs, we visualized the abundance and localization of viral RNA in infected AMs or IMs by confocal and super-resolution (SR) microscopy using smFISH probes tiled against positive strand viral RNA (Orf1a and N genes), and antibodies against dsRNA (replication intermediate) and the lysosomal marker LAMP1. Based on expression of dsRNA and viral gRNA, as well as aspects of the nuclear, lysosomal, and overall morphology of the cell, we distinguished five phenotypic classes among the infected macrophages (Fig. 5b-c, Classes I - V, Methods); we then mapped these major cellular events onto the known progression of the coronavirus life cycle, beginning with transcription of dsRNA intermediates, followed by replication of the full length genomic RNA, and packaging and release of the matured virions^61,62^ (Classes I - III, early, intermediate, and advanced viral replication; Class IV, viral aggregates; Class V, host cell destruction and death).

The early stages of viral infection appear to be comparable or even more efficient in AMs than IMs, as indicated by the abundance of Class I (early viral replication, expressing only dsRNA) intermediates observed with untreated virions (AMs 6%, IMs 6%), and with UV-inactivated virions (AMs 10.3%, IMs 2.6%) that do not proceed beyond this stage (Fig. 5d). But viral replication and accumulation is more efficient in IMs than AMs, as indicated by the greater abundance of Class II (intermediate viral replication) intermediates expressing both dsRNA and viral gRNA, presumably beginning to replicate the full length genomic RNA (AMs 1.6%, IMs 4.6%), and Class III (advanced viral replication) intermediates that have progressed to expressing exclusively viral gRNA and generally at high abundance (AMs 0.9%, IMs 5.8%). Together, the relative abundance of these Class II and III ‘late viral replicating’ cells (AMs 2.5%, IMs 10.4%) is consistent with the plaque assay results demonstrating higher virion production in IMs than AMs (Fig. 5f).

The most striking difference between infection of IMs and AMs was that only infection of IMs resulted in formation of large (>500 um) and globular aggregates of viral gRNA that consumed the cytoplasm, what we term viral “RNA bodies” (Class IV, AMs 0%, IMs 3.7%, Fig. 5c,d). Super-resolution (SR) examination of these viral RNA bodies by single molecular localization microscopy confirmed their structure as dense without finer localizations (Fig. 5e), contrasting with the thousand-fold smaller (<0.25 um) and punctate perinuclear gRNA localizations in Classes II/III (Fig. 5e), and with previous SR characterizations of coronavirus infection^63^. Finally, the terminal stage of IM infection was the obliteration of nuclear and/or lysosomal architecture and overall cellular morphology, with abundant but diffuse viral gRNA (Class V, AMs 0%, IMs 2.2%, Fig. 5c), whose beginnings were apparent in Class IV with nuclear blebbing and lysosomal compaction around the nucleus instead of a smooth perinuclear ring. These late or terminally-infected phenotypes likely correspond to the IMs that have been taken over by SARS-CoV-2, leading to their ultimate demise and death.

### Viral entry into interstitial macrophages uses DC-SIGN/CD209 but not ACE2

To explore the mechanism of SARS-CoV-2 entry into lung macrophages, we developed a SARS-CoV-2 Spike-pseudotyped lentivirus (lenti-S-NLuc-tdT) encoding nanoluciferase (NLuc) bioluminescence and tdTomato fluorescence dual readout reporters (Fig. 5g, scheme). We tested whether this pseudotyped lentivirus could enter AMs or IMs by exposing freshly purified human primary AMs or IMs to the virus for 48 hours in culture and then assaying NLuc luminescence. Whereas a control lentivirus lacking SARS-CoV-2 Spike protein did not elicit measurable luminescence in either lung macrophage population, the SARS-CoV-2 Spike-pseudotyped lentivirus elicited robust NLuc luminescence in both AMs and IMs (Fig. 5g). Thus, SARS-Cov2 Spike protein mediates entry into both types of human lung macrophages.

Neither lung macrophage population expressed detectable mRNA levels of *ACE2*, the canonical SARS-CoV-2 receptor, by scRNA-seq (Extended Data Fig. 11) or smFISH (Fig. 2c). However, examination of the expression of other proposed SARS-CoV-2 receptors (Extended Data Fig. 10a) and host factors (Extended Data Fig. 10b) revealed differential expression of a family of lectin proteins that have been newly identified and proposed as SARS-CoV-2 attachment receptors^64,65^: *DC-SIGN* (CD209), an N-glycan-binding C-type lectin, was selectively expressed only in activated IMs, whereas *SIGLEC1* (CD169), a sialic-acid-binding immunoglobulin-like lectin, was selectively expressed only in AMs. To test the function of these candidate host cell receptors in viral entry, we examined the effect of blocking antibodies against them in the pseudotyped lentivirus assay described above. Blocking antibodies against DC-SIGN/CD209 inhibited lenti-S-NLuc-tdT entry into IMs (50% neutralizing titer, NT50: 0.32 μg/ml), whereas blocking antibodies against Siglec-1/CD169 had no effect (Fig. 5h). We found just the opposite for AMs: blocking antibodies against Siglec-1/CD169 (NT50: 0.02 μg/ml) inhibited lenti-S-NLuc-tdT entry, whereas blocking antibodies against DC-SIGN/CD209 had no effect. Interestingly, although blocking antibodies against ACE2 had no inhibitory effect on lenti-S-NLuc-tdT entry into IMs as expected from their lack of ACE2 expression (Extended Data Fig. 10a, Extended Data Fig. 11), ACE2 blocking antibodies did reduce entry into AMs (NT50: 2.75 μg/ml), which despite the lack of detectable *ACE2* mRNA (Extended Data Fig. 10a) nevertheless expressed ACE2 on their surface (Extended Data Fig. 11). We also tested two clinically relevant antibodies that hinder ACE2 binding (COVA2-15, REGN-CoV) and found that although both reduced viral entry into AMs, neither affected entry into IMs (Extended Data Fig. 12).

Thus, while SAR-CoV-2-Spike dependent entry into alveolar macrophages depends on SIGLEC-1/CD169 and the canonical receptor ACE2, entry into interstitial macrophages uses a DC-SIGN/CD209 mechanism independent of ACE2.

## DISCUSSION

We established an experimental model of COVID-19 initiation in the human lung by productive infection of *ex vivo* cultured human lung slices with SARS-CoV-2. scRNA-seq and smFISH generated a comprehensive atlas of SARS-CoV-2 lung cell tropism and allowed us to probe the viral life cycle and its dynamic effects on the corresponding host gene expression program of individual lung cell types, by reconstruction of “infection pseudotime” from the single cell profiles of infected intermediates. The results indicate that the most susceptible lung target of SARS-CoV-2 and initial focus of inflammation and fibrosis is activated interstitial macrophages. In this newly characterized lung macrophage subtype, viral RNA amplification results in host cell takeover with viral transcripts comprising up to 60% of the total cellular transcriptome. During takeover, there is cell-autonomous induction of an interferon-dominated inflammatory response, including induction of five chemokines that can recruit local innate immune cells expressing the cognate receptors, including DCs (via *CCL2*, *CCL13, CXCL10*), neutrophils (*CCL8, CCl13*), and additional macrophages (*CCL8, CCL13*) forming an autocatalytic cycle, as well as CD4 and CD8 T cells (*CXCL10*). Takeover also induces expression of pro-inflammatory cytokine *IL6*, among the molecules critical to cytokine storm^60^, which can signal to many immune cells and most structural cells of the lung, as well as the pro-fibrotic cytokine *TGFB1*, the central mediator of lung fibrogenesis^66^, which can signal to most epithelial and fibroblast cell types. Thus, SARS-CoV-2 infection and takeover of interstitial macrophages and interferon-dominated induction of this suite of chemokines and cytokines forms an early focus of lung inflammation, immune infiltration, and fibrosis.

While our studies show that macrophages are the most prominent cell targets in the human lung, they also reveal that two distinct molecular lineages of macrophage targets, interstitial (IMs) and alveolar (AMs) macrophages, each patrolling a different lung compartment, respond completely differently to SARS-CoV-2 (Fig. 6e), with important implications for its pathogenesis mechanism and therapeutics. Although both purified AMs and IMs supported production of infectious virions, the virus enters the cells by distinct mechanisms: infection of AMs uses the canonical receptor ACE2 and CD169, whereas infection of IMs uses an ACE2-independent mechanism that relies instead on CD209. And, while the initial stage of viral replication with formation of dsRNA intermediates is similar or more efficient in AMs, later stages of replication and production of infectious virions is greater in IMs. Finally, whereas viral replication and induction of an innate immune response is muted in AMs, the virus takes over IMs, dominating the host cell transcriptome and forming large globular viral RNA bodies that appear throughout the cytoplasm, triggering or amplifying the potent inflammatory and fibrotic response, as well as activating inflammasomes, ultimately obliterating cell architecture and destroying the host cell. The results with the purified macrophage cell types reveal the intrinsic differences between their infection cycles, and they suggest that while AMs are competent to restrict viral amplification without an obvious transcriptional response, the robust response to the virus in IMs occurs too late or lacks some critical component(s) of an effective antiviral response, allowing the viral RNA to accumulate, thereby amplifying the inflammatory response, forming viral RNA bodies, and ultimately destroying the host cell. It will be important to determine in future studies if the different entry receptors, innate antiviral competence, and/or other distinguishing features of the host cell environment determine the distinct infection cycles in IMs and AMs.

Although prior studies of viral RNA or protein accumulation have begun to implicate myeloid cells in SARS-CoV-2 infection^17,21,22,24,67^, none distinguished the two lung macrophage types or suggested production of infectious virions. Recent studies in fact indicate just the opposite: no infection of macrophages^68^ and only abortive infection of monocytes^23^. The SARS-CoV-2 viral life cycle thus differs dramatically not only between lung resident IMs and AMs, but also among closely related lineages or cell lines (such as monocytes and bone-marrow-derived macrophages). This sensitive dependence of the outcome of viral infection on host cell type highlights the need in future COVID-19 studies to resolve the molecular subtype of lung macrophage infected, the extent of viral takeover, and the specific host cell response, in evaluating their role in the disease.

Based on our data we propose an initial model in which the two types of lung macrophages have different contributions to COVID-19 pathogenesis (Fig. 6). AMs patrolling the airspace are expected to be among the first cells encountered by virions reaching the alveoli. Infection of AMs results in viral replication and production at low levels, evading induction of a host immune response but presumably contributing to interalveolar spread of the virus. Infection of epithelial cells lining the alveolus, most prominently alveolar type 2 (AT2) cells can lead to high viral RNA levels that presumably alter and destroy them, injuring the alveolar epithelium and compromising its repair and its barrier function (Fig. 6a). Through transepithelial spread or an epithelial breach, IMs become infected, further propagating the virus interstitially and initiating viral pneumonia (Fig. 6b). Viral takeover of IMs then triggers the inflammatory and fibrotic phase of COVID-19 by induction of a specific suite of cytokines and chemokines (Fig. 6c), explaining the main pathologies observed in COVID-19 ARDS: local immune recruitment, activation, and infiltration, as well as extensive interstitial fibrosis, resulting in the respiratory demise (hypoxemic respiratory failure) and pathology (diffuse alveolar damage) characteristic of COVID-19 ARDS^4,69^. Breakdown of the endothelial barrier could facilitate spread of the virus and release of IL6 and other lung inflammatory signals into the bloodstream, commencing the systemic effects of cytokine storm^60^ (Fig. 6d).

The central role in the model of infected interstitial macrophages in the transition of COVID-19 pneumonia to ARDS and cytokine storm implies that blocking their infection would prevent the most serious consequences of COVID-19. In this regard, our data showing that IM entry uses an ACE2-independent mechanism that relies on CD209 and a region of the viral Spike-protein outside the canonical ACE2 interface may explain why therapeutic antibodies have invariably failed in severe cases of COVID-19 lung inflammation – they block viral entry into airway and alveolar epithelial cells that initiate the disease, but not the interstitial macrophages we propose catalyze the inflammatory and fibrotic phase (Fig. 6e). Effective therapies to prevent the onset or reverse the progression of severe COVID-19 ARDS should address both the noncanonical pathway of viral entry and the molecular and cellular consequences of the downstream inflammatory and fibrotic cycles.

Our approach for elucidating the molecular and cellular basis of COVID-19 initiation relied on: productive infection of a human lung slice system and scRNA-seq pipeline that allowed culture, capture, and gene expression analysis of both viral and host transcriptomes in comprehensive cell types of the native organ; careful comparison with freshly-harvested tissue to distinguish direct virus-induced changes from culture-induced effects; and development of the computational method “infection pseudotime” to reveal the early cell-intrinsic gene expression program induced by viral infection. Although the lung slices deployed here contain a nearly complete complement of human lung cells and interactions, they do not have the full native geometry or physiology of the human lung. Future improvements should focus on developing such fully native models of the human lung, complete with recruitment of circulating immune cells, to directly test and extend the pathogenesis model proposed above based on results with the slice cultures and purified cells and virions. Future studies should also seek to improve the resolution of spatial patterns of gene expression (including mRNA as well as protein expression) and cellular structures, interactions, and behaviors, as well as characterization of viral subgenomic transcripts to resolve gene-level viral expression in single cells. Our approach can be used to elucidate the initiation program and evaluate therapeutics for any human lung infection at cellular resolution.

## METHODS

### *Ex vivo* culture of human lung tissue

Fresh, normal lung tissue was procured from organ donors that have exhausted therapeutic recipient options through Donor Network West, or intraoperatively from surgical resections at Stanford Hospital. Case 1 was a male organ donor aged 62. The entire lung was obtained en bloc, and the left upper lobe (LUL) was selected based on clear imaging as indicated on the donor report. Case 2 was a female organ donor aged 36 with a history of M to F gender reassignment. The entire left lung was obtained, and the LUL was selected based on clear imaging as indicated on the donor report. Case 3 was a 57-year-old-female with a history of fatty liver disease, diagnosed with stage two adenocarcinoma, who underwent left lower lobe (LLL) lobectomy. Case 4 was an 83-year-old female with a distant smoking history, diagnosed with stage two adenocarcinoma, who underwent LUL lobectomy. Case 5 was a male organ donor aged 57. The entire lung was obtained en bloc, and small sections were cut from RUL, RML, and RLL, based on clear imaging as indicated on the donor report. Case summaries are provided in Table S4.

In each case, proximal (airway) and distal (alveolar) regions were resected and cut into 300-500 µm slices with platinum coated double edge blade (Electron Microscopy Sciences 7200301) manually. For donor lungs, the healthiest lobe was selected based on absence of inflammation or infection detected by qPCR or chest imaging in the donor summary. Both airway and alveolar slices (3 or 4 total) were cultured in the same well in a 12-well plate with or without precoating of 500 µL of growth factor reduced Matrigel (Corning 354230), 1 mL DMEM/F12 media supplemented with GlutaMAX (Gibco 35050061) 10% fetal bovine serum (FBS, Gibco 10-082-147), 100 U/mL Penicillin-Streptomycin (Gibco 15140122), and 10 mM HEPES (N-2-hydroxyethylpiperazine-N-2-ethane sulfonic acid, Gibco 15630080), and incubated at 37°C, 5% CO_2_.

Tissue samples obtained from surgical resections were obtained under a protocol approved by the Stanford University Human Subjects Research Compliance Office (IRB 15166) and informed consent was obtained from each patient before surgery. All experiments followed applicable regulations and guidelines.

The research protocol for donor samples was approved by the DNW’s internal ethics committee (Research project STAN-19-104) and the medical advisory board, as well as by the Institutional Review Board at Stanford University which determined that this project does not meet the definition of human subject research as defined in federal regulations 45 CFR 46.102 or 21 CFR 50.3.

### Cell lines

VeroE6 cells were obtained from ATCC as mycoplasma-free stocks and maintained in supplemented DMEM (DMEM (Dulbecco’s Modified Eagle Medium) (Thermo 11885-092) with 1X L-glut (Thermo SH30034), MEM nonessential amino acids (Thermo 11140050), 10mM HEPES (Gibco 15630-080), 1X antibiotic/antimycotic (Life Technologies SV30079), and 1mM sodium pyruvate (Gibco 11360-070)) with 10% heat-inactivated (HI) FBS (Sigma F0926). Vero/hSLAM cells were a kind gift from Dr. Chris Miller and Dr. Timothy Carroll (University of California, Davis) and were mycoplasma free (PlasmoTest, Invivogen). VeroE6/TMPRSS2 cells^70^ were obtained from the Japanese Collection of Research Bioresources Cell Bank as mycoplasma-free stocks. Vero/hSLAM and VeroE6/TMPRSS2 were maintained in supplemented DMEM with 10% HI-FBS and 1 mg/mL G418 sulfate (Thermo Fisher 10-131-027). HeLa/ACE2/TMPRSS2 cells were a generous gift from Dr. Jesse Bloom at the Fred Hutchinson Cancer Research Center.

### SARS-CoV-2 infections

#### Preparation of viral stock

SARS-CoV-2 (USA-WA1/2020) was obtained in March 2020 from BEI Resources and passaged in VeroE6 cells in supplemented DMEM with 2% HI-FBS. Viral stocks were cleared of cellular debris by centrifugation (1,000*g,* 10 min, 4℃), aliquoted, and stored at -80℃. Titer was determined by plaque assay (see below). The viral stock was verified by deep sequencing (∼100,000X coverage per base) against the reference sequence (GenBank MN985325.1), and all tissue replicates were infected with passage 3 virus. A purified stock (“WA1 new”) was also made by passaging in Vero/hSLAM cells, then clarifying by centrifugation (4,000*g*, 10 min, 4℃) followed by three buffer exchanges of phosphate-buffered saline (PBS) using Amicon Ultra-15 100 kDa Centrifugal Filter Units (Millipore Sigma). This viral stock was also verified by deep sequencing.

#### Lung slice infections

Infections of lung slices were performed in supplemented DMEM with 2% FBS at a multiplicity of infection (MOI) of 1 (assuming all lung cells in the culture could be target cells) at 37℃. After 2 hours, free virions were removed by washing the tissue with PBS, after which the slices were cultured in DMEM/F12 with 10% FBS for 24 or 72 hrs. All procedures involving infectious SARS-CoV-2 were performed inside a class II biosafety cabinet (BSC) in the CDC-approved Biosafety Level 3 (BSL3) facility at Stanford University under approved biosafety protocols. All stocks generated and used were between 0.5 and 2 x 10^6^ pfu/mL.

#### Purified macrophage infections

At least 80,000 - 100,000 purified AMs or IMs (see below) were plated in µ-Slide 8 Well chamber slides (Ibidi) and cultured in macrophage culture medium (DMEM/F12 media supplemented with 10% FBS, 100 U/mL Penicillin-Streptomycin, and 50 ng/ml M-CSF), and incubated at 37°C, 5% CO_2_, for 1 – 3 days to allow cells to rest. Infections of macrophages were performed in supplemented DMEM with 2% FBS at the indicated multiplicity of infection (MOI) at 37℃. After 2 hours, cells were gently washed with Dulbecco’s PBS with calcium and magnesium (Gibco), followed by replacement with macrophage culture medium.

#### Plaque assay

VeroE6 or VeroE6/TMPRSS2 cells were plated at 4.5-5 x 10^5^ cells/well in a standard 12-well tissue culture plate (Falcon) one day prior to infection. On the day of infection, cells were washed once with PBS. 100 μL of lung slice culture supernatants were added to the monolayer undiluted or diluted as indicated in supplemented DMEM containing 2% FBS. After 45 min of rocking inside an incubator at 37℃ to allow viral adsorption to the cells, plates were overlaid in the BSC with a fresh, pre-warmed (37℃) 1:1 mixture of 2X MEM (Thermo 11935046) (supplemented with 0.24% bovine Serum Albumin (BSA, Sigma A9576), 2X L-glutamate, 20mM HEPES, 2X antibiotic/antimycotic (Life Technologies), and 0.24% sodium bicarbonate (Sigma S8761)) and 2.4% Avicel (FMC Biopolymer). Plates were then returned to the incubator for 72h (VeroE6) or 48h (VeroE6/TMPRSS2) prior to overlay removal, washing with PBS, fixation with 70% ethanol, and staining with 0.3% crystal violet (Sigma). For the timecourse (Extended Data Fig.1), lung slices were infected, washed, and placed in culture media. At 24h the supernatant was harvested, stored frozen, and replaced completely with fresh media. At 72h, the supernatant was harvested and stored frozen. The supernatants were then thawed, and plaque assays performed on the same plate as above.

#### Viral inactivation and remdesivir treatment

UV inactivation of virus was performed by delivering 1800 MJ of UV-C light (254 nm) to 250 uL of undiluted viral stock in a 24-well plate using a Stratalinker 1800 (Stratagene California, La Jolla, CA) inside a BSC in the BSL3. For heat inactivation, one aliquot of thawed undiluted viral stock was placed in a heat block at 60℃ for 20 minutes inside a BSC in the BSL3. Inactivations were verified by plaque assay. For remdesivir (RDV) treatment, 10 mM stocks of RDV (Gilead) in DMSO were prepared and added to lung slices cultures at the time of infection to a final concentration of 10 µM. Slices were re-dosed after washing off the virus inoculum.

### Single-cell mRNA sequencing

#### Lung cell isolation

All fresh (non-cultured and non-infected) tissue was processed at BSL2, and all cultured or infected tissue was processed in BSL3.

BSL2: Normal lung tissue was obtained as described for the slice cultures. All tissues were received and immediately placed in cold PBS and transported on ice directly to the research lab. Individual human lung samples were dissected, minced, and placed in digestion media (400 μg/ mL Liberase DL (Sigma 5466202001) and 100 μg/mL elastase (Worthington LS006365) in RPMI (Gibco 72400120) in a gentleMACS c-tube (Miltenyi 130-096-334). Samples were partially dissociated by running ‘m_lung_01_01’ on a gentleMACS Dissociator (Miltenyi 130-093-235), incubated on a Nutator at 37 °C for 30 min, and then dispersed to a single cell suspension by running ‘m_lung_02_01’. Processing buffer (5% FBS in 1XPBS) and DNase I (100 μg/mL, Worthington LS006344) were then added, and the samples rocked at 37 °C for 5 min. Samples were then placed at 4 °C for the remainder of the protocol. Cells were filtered through a 100-μm filter (Fisher 08-771-19), pelleted (300*g*, 5 min, 4 °C), and resuspended in ACK red blood cell lysis buffer (Gibco A1049201) for 3 min, after which the lysis buffer was inactivated by adding excess processing buffer. Cells were then filtered through a 70-μm strainer (Fisher 22363548), pelleted again (300*g*, 5 min, 4 °C), and resuspended in 2% FBS in PBS.

BSL3: After washing off virus and incubating for the indicated times, lung slices were washed with PBS and carefully transferred to 15 mL conical tubes (Falcon) containing 5 mL digestion buffer (DMEM/F12 with 400 µg/mL Liberase DL (Sigma), 50 µg/mL elastase, and 250U benzonase (EMD Millipore 706643)) and incubated with manual or automatic rocking at 37℃ for 1 hour, followed by serum neutralization of Liberase and elastase activity with 10% FBS in cold DMEM/F12 media. For infection 1 only, the tissue was then dissociated by running m_lung_02 on a gentleMACS dissociator inside the BSC. The tissue was then mashed through a 100 µM filter with a syringe insert (Falcon), and the filter was washed with additional cold DMEM/F12 with 10% FBS to recover any remaining cells. The cellular suspension was spun at 4℃ at 300 x *g* for 5 minutes, washed, and exposed to 1mL cold ACK lysis buffer (Sigma) for 1 minute on ice. The lysis buffer was neutralized by dilution with 5 mL cold DMEM/F12 with 10% FBS, after which the cells were pelleted and resuspended in DMEM/F12 with 10% FBS, and the cells were stained with Trypan blue (Sigma T8154), sealed out of the BSC, and counted manually. For all steps, cells were kept at 0-4℃ using a cold block (Eppendorf Isotherm system).

#### 10x mRNA capture, library construction, and sequencing

BSL2: Cells isolated from normal lung tissue or purified by FACS were captured in droplet emulsions using a 10x Chromium Controller (10x Genomics). cDNA was recovered and libraries were prepared using 10x Genomics 3’ or 10x Genomics 5’ Single Cell V3.1 protocol (infections 1,2,4 and 5 were sequenced using 3’ chemistry, while infection 3 used both 3’ and 5’ technology), as described^34^. Sequencing libraries were analyzed (Agilent TapeStation D4150, using regular and high sensitivity D5000 ScreenTape) and balanced, and sequenced on a NovaSeq 6000 (Illumina).

BSL3: The 10x Genomics Single Cell protocols were performed as before, with the following modifications for BSL3. The 10x Genomics, 3’ or 5’ Single Cell v3.1 master mix was prepared outside the BSC. Within the BSC, cells prepared as above were added to the master mix in PCR tubes (USA Scientific 1402-4708) in a 96-well cold block (Ergo 4ti-0396) and the 10x chip was loaded per manufacturer’s instructions, sealed, and processed in a 10x Chromium Controller in the BSC. The resultant cell/bead emulsions were loaded into PCR tubes and transferred immediately to a pre-warmed (53℃) PCR machine for cDNA synthesis carried out at 53℃ for 45 minutes, then 85℃ for 5 minutes, then 60℃ for 15 minutes (plaque assays showed that exposure of SARS-CoV-2-infected samples at 60℃ for 20 minutes in this manner rendered the sample non-infectious). After cDNA synthesis, samples were transferred out of the BSL3 for cDNA recovery, amplification, and sequencing library preparation as above.

#### Sequencing read alignment

Sequencing reads from single cells isolated using 10x Chromium were demultiplexed and then aligned to a custom built Human GRCh38 (GENCODE v30) and SARS-CoV-2 WA1 (GenBank: MN985325.1) reference using Cell Ranger (version 5.0, 10x Genomics).

#### Iterative cell clustering and annotation

Expression profiles of cells from different subjects were clustered separately using Python software package Scanpy (v1.7.2). For host genes, normalization was performed as described^34^;Unique molecular identifiers (UMIs) were normalized across cells, scaled per 10,000 using the “sc.pp.normalize_total” function, converted to log-scale using the “sc.pp.log1p” function, and highly variable genes were selected with the “sc.pp.highly_variable_genes” function with a dispersion cutoff of 0.5, and scaled to unit variance and zero mean (z-scores) with the “scanpy.pp.scale” function, clipping values exceeding standard deviation 10. Principal components were calculated for these selected genes with the “sc.tl.pca” function. Clusters of similar cells were detected using the Leiden method (“tl.leiden” function) for community detection including only biologically meaningful principle components, as described^34^, to construct the shared nearest neighbor map (“sc.pp.neighbors”) and an empirically set resolution, visualized by uniform manifold approximation and projection (UMAP; “tl.umap” function).

Cells were iteratively clustered as described^34^, with the following modifications. After separating clusters by expression of tissue compartment markers, cultured cell types generally segregated from their non-cultured counterparts. When possible, we assigned cell types to the canonical cell types using the most sensitive and specific markers identified in the human lung cell atlas^1^. For culture-induced subtypes that showed substantial transcriptional change, a representative marker gene was prepended to their canonical identity (e.g., IRF1+ aCap). If the transcriptional change caused the cell type to lose markers that define their canonical identity, we named them based on the general type that could be assigned, and prepended a representative marker gene (e.g., KLF6+ Endo). If most of the cluster-specific markers were ribosomal or mitochondrial genes, they were labeled as low quality (e.g., Stromal-LQ). If most of the expressed genes were viral and we could not distinguish which cell type the cluster belonged to due to downregulation of marker genes, they were designated “infected” (e.g., Infected-LQ). Cells from different subjects with the same annotation were merged into a single group for all downstream analyses. Cell types that were exclusively found to be culture induced were grouped as “culture induced” (e.g., Induced Fibroblast) for viral tropism analysis.

Some native subtypes characterized by subtle transcriptional differences could not be resolved by droplet-based 10x sequencing (e.g., proximal subtypes for basal or ciliary cells, molecular subtypes of bronchial vessel cells, mast/basophils), and several rare (neuroendocrine cells, ionocytes) or anatomically-restricted cell types (e.g. serous cells in submucosal glands) were absent from the profiled lung tissue.

#### Viral takeover analysis

For the UMIs that aligned to the SARS-CoV-2 viral genome, raw UMIs were either directly converted to log scale (“log10(Viral UMIs + 1)”) or explicitly divided by total cellular UMIs but not log-converted (“Viral UMIs”). Viral takeover trends were visualized by non-parametric Local Regression (LOESS, R stats version 3.6.2).

#### Viral pseudotime analysis

For viral pseudotime analysis, computations were performed in R using the Seurat package (v3). Infected alveolar macrophages (AMs) and activated interstitial macrophages (a-IMs) from infection 1 were grouped, and counts were normalized using the ‘SCTransform’ command. Principal component analysis was performed using the ‘RunPCA’ command with default parameters and visualized with ‘DimHeatmap’. To identify the major axes of variation within the infected macrophage subtypes that best correlated with SARS-CoV-2 RNA levels, the principal components with significant contribution from SARS-CoV-2 counts (among the top 15 genes with highest loadings) were selected for further inspection. PC.1 was found to be associated with increasing viral RNA levels in both AMs and a-IMs, and PC.2, PC.3, and PC.4 were found to be associated with increasing viral RNA levels only in a-IMs. To isolate the genes that specifically associated with increasing SARS-CoV-2 viral RNA levels in AMs, PC.3 was subtracted from PC.1 (see Extended Data Fig.5).

Thus, infection pseudotime was defined respectively for AMs and a-IMs as progression along the following axes by taking the following linear combinations of principal components:

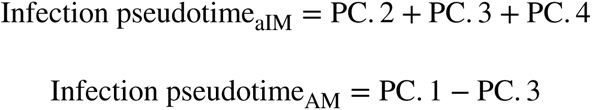

AMs and aIMs were assigned respective pseudotime values that were normalized between 0 and 1.

#### Subgenomic RNA analysis

To detect viral subgenomic RNA junctions, we ran SICILIAN^43^, a statistical wrapper that takes alignment files from a spliced aligner and calls true positive RNA splice junctions by employing a statistical model. SICILIAN assigns an empirical p-value to each detected junction in a 10x dataset, quantifying the statistical confidence of each detected junction being a truly expressed RNA junction. We used STAR v.2.7.5a as the aligner and aligned fastq files from all infections to our custom built Human GRCh38 (GENCODE v29) and SARS-CoV-2 WA1 (GenBank: MN985325.1) reference. STAR was run in two-pass mode, in which the first pass identifies novel splice junctions and the second pass aligns reads after rebuilding the genome index with the novel junctions and its parameters were tuned to avoid bias against non-GTAG junctions as previously shown^71^.

#### Immunostaining and single molecule in situ hybridization

BSL2: Samples were fixed in either 10% neutral buffered formalin, dehydrated with ethanol and embedded in paraffin wax, as described^34^.

BSL3: Slices not taken for digestion were washed with PBS and transferred to 15 mL conical tubes containing 10% neutral buffered formalin (Sigma) and held at 4°C for 72 hours prior to transfer out of the BSL3. Slices were then transferred to 15 mL conical tubes containing PBS prior to dehydration.

Sections (6 μm) from paraffin blocks were processed using standard pre-treatment conditions for each per the RNAscope multiplex fluorescent reagent kit version 2 (V2) Assay (Advanced Cell Diagnostics, (ACD)), or immunostaining RNAscope co-detection assay in which antibody labeling was carried out after RNAscope V2 assay, or RNAscope HiPlex Assay protocols. AlexaFluor plus secondary antibodies (488 plus, anti-mouse, Invitrogen A32723; 750, anti-rabbit, Invitrogen A21039) were used at 1:1,000 dilution. For RNAscope V2 assays, TSA-plus fluorescein, Cy3 and Cy5 fluorophores were used at 1:500. Micrographs were acquired with laser scanning confocal fluorescence microscopy (Leica Stellaris 8) and processed with ImageJ and Imaris (version 9.2.0, Oxford Instruments). smFISH experiments were performed on lung tissue from at least two human participants distinct from the donors used for sequencing, and quantifications were based on at least 10 fields of view in each. For smFISH, fields of view were scored manually, calling a cell positive for each gene probed if its nucleus had at least two associated expression puncta.

The following primary antibodies were used at 1:100: CD68 (mouse, Abcam ab955), RAGE (rabbit, Abcam ab216329), ASC (mouse Sigma 04-147). The following V2 RNAscope probes were used: *MSR1* (ACD 468661), *RNASE1* (ACD 556551), *FABP4* (ACD 470641), *IER3* (ACD 1000371), *nCoV2019-S* (ACD 845701); the following HiPlex probes were used: *ACE2* (ACD 848151), *DPP4* (ACD 477549), *EPCAM* (ACD 310288), *COL1A2* (ACD 432721), *PTPRC* (ACD 601998), *ASPN* (ACD 404481), *nCoV2019-S* (ACD 848561), *nCoV2019-orf1ab-sense* (ACD 859151), *CLDN* (ACD 517141), *EDNRB* (ACD 528301), *AGER* (ACD 470121), *SFTPC* (ACD 452561), *CD4* (ACD 605601), *CD3-pool* (ACD 426621), *CD8A* (ACD 560391), *MARCO* (ACD 512231), *STAB1* (ACD 472161), *FABP4* (ACD 470641), *FOXP3* (ACD 418471), *IER3* (ACD 1000371).

#### Macrophage isolation and enrichment

Lung tissue was obtained as described above for the slice cultures. All tissues were received and immediately placed in cold PBS and transported on ice directly to the research lab. Individual human lung samples were dissected, minced, and placed in digestion media (400 μg/mL Liberase DL (Sigma 5466202001) and 100 μg/mL elastase (Worthington LS006365) in RPMI (Gibco 72400120) in a gentleMACS c-tube (Miltenyi 130-096-334). Samples were partially dissociated by running ‘m_lung_01_01’ on a gentleMACS Dissociator (Miltenyi 130-093-235), incubated at 37 °C for 30 min, and then dispersed to a single cell suspension by running ‘m_lung_02_01’.

Processing buffer (5% fetal bovine serum in PBS) and DNase I (100 μg/mL, Worthington LS006344) were then added, and the samples rocked at 37 °C for 5 min. Samples were then placed at 4 °C for the remainder of the protocol. Cells were filtered through a 100-μm filter (supplier), pelleted (300g, 5 min, 4 °C), and resuspended in ACK red blood cell lysis buffer (Gibco A1049201) for 3 min, after which the buffer was inactivated by adding excess processing buffer. Cells were then filtered through a 70-μm strainer (Fisherbrand 22363548), pelleted again (300g, 5 min, 4 °C), and resuspended in magnetic activated cell sorting (MACS) buffer (0.5% BSA, 2 mM EDTA in PBS) with Human FcR Blocking Reagent (Miltenyi 130-059-901) to block non-specific binding of antibodies. The isolated lung cells were stained with CD206 antibodies conjugated to biotin (Miltenyi 130-095-214), washed twice with MACS buffer, then stained with Anti-Biotin MicroBeads (Miltenyi 130-090-485) and passed through an LS MACS column on a MidiMACS Separator magnet or a SuperMACS II Separator magnet, with the retained population designated “MACS CD206^+^” and further purified by FACS (see below).

#### Fluorescence activated cell sorting (FACS) purification of AMs and IMs

The MACS CD206^+^ enriched population of lung resident macrophages were incubated with FcR Block (Biolegend 422302) for 5 minutes and stained at a dilution of 1:50 with one of two panels of directly conjugated antibodies for 30 minutes at 4 °C: anti-human CD45 (BD 563792), CD204 (Biolegend 371906), CD206 (Biolegend 321132), CD14 (BD 562698), CD16 (BioLegend 302028), ACE2 (R&D FAB933P), HLA-DR (BioLegend 307618), CD11b (BioLegend 393114), CD11c (BioLegend 301644); anti-human CD45 (BioLegend 324016), CD204 (BioLegend 371904), and CD206 (BioLegend 321103). Stained cells were then washed with FACS buffer (2% FBS in PBS) three times, then incubated with cell viability marker propidium iodide (PI, 1 μg/mL, Biolegend 421301). Flow cytometry was performed on a FACS Aria II (BD Biosciences).

Living (PI^-^) single, immune (CD45^+^), lung resident macrophages (CD206^+^) were stained for the above panel of cell surface antigens that have previously been suggested to segregate them into alveolar macrophages (AMs) and interstitial macrophages (IMs), and that are differentially expressed according to the scRNA-seq transcriptomic profiles obtained from lung slice culture, and sorted into CD206^+^CD204^hi^ and CD206^+^CD204^lo^ populations. The sorted populations were directly subjected to 10x single cell mRNA sequencing at BSL2 as described above, which confirmed their molecular identities as AMs and IMs, respectively.

#### RNA hybridization chain reaction (HCR) and immunofluorescence (IF) staining

Purified AMs or IMs cultured on slides were infected with untreated (viable) or UV-C inactivated SARS-CoV-2 virions, then fixed in BSL3 by transfer of the slides into 15 mL conical tubes containing 10% neutral buffered formalin (Sigma) and held at 4°C for 72 hours prior to transfer out of the BSL3. Slides were then transferred to 15 mL conical tubes containing PBS, prior to cell permeabilization with 70% ethanol overnight at -20°C. After decanting the ethanol, cells were washed once with PBS at room temperature for 5 min. Slides were blocked with antibody buffer (Molecular Instruments) for 1 hour, followed by incubation with primary antibodies at a dilution of 1:100 overnight (>12 h) at 4°C, washed three times with 1x PBST (0.1% Tween-20), then incubated with secondary antibodies at a dilution of 1:1,000 for 1 h at room temperature, and washed three times with 1x PBST before post-fixing with 4% paraformaldehyde for 10 mins at room temperature.

RNA hybridization chain reaction was carried out according to manufacturer instructions (Molecular Instruments). Briefly, RNA hybridization was performed by immersing the immunostained slides in 5x SSCT (SSC with 0.1% Tween-20) for 5 min at 37°C, then hybridized with 16 nM RNA HCR probe solution overnight (>12h) at 37°C in a humidified chamber, and excess probes washed off using a 25%–100% 5xSSCT step gradient (at 25% steps) with 15 minutes for each incubation. RNA amplification was performed using 6 pmol of hairpin h1 and 6 pmol of hairpin h2 that were separately snap-cooled (by heating to 95°C for 90 seconds then cooled to room temperature in a dark drawer for 30 min). The cooled hairpins were then mixed to create a 60 nM hairpin solution, which was used to incubate slides overnight (>12 h) in a dark humidified chamber at room temperature. Excess hairpin was removed by washing three times with 5x SSCT, and slides were then counterstained with DAPI and mounted for laser scanning confocal fluorescence imaging (Stellaris 8), or super-resolution imaging (see below).

#### Analysis and scoring of infected macrophages

Purified, infected, and stained AMs or IMs were quantified by confocal microscopy and analyzed in Imaris (version 9.2.0, Oxford Instruments), and resolved six phenotypic classes based on relative expression of dsRNA and viral gRNA, as well as aspects of the nuclear, lysosomal, and overall morphology of the cell. Class 0 cells express neither dsRNA nor viral gRNA, hence were not infected; they comprised the vast majority of cells (91% of AMs and 78% of IMs). Class I cells only express dsRNA, are observed rarely in the UV-inactivated viron conditions, and in the infection conditions comprised 6% of AMs and 3.4% of IMs; these are inferred to be the earliest stage of viral infection, representing entry of input virions or the very earliest stages of viral replication. Class II cells express both dsRNA and viral gRNA, presumably reflecting the replication of full length genomic RNA, and comprised 1.6% of Ams and 4.6% of IMs. Class III cells exclusively express viral gRNA, generally at high abundance, and comprised 0.9% of AMs and 5.8% of IMs. Although there were more Class I (Early) AMs, the Class II and Class III cells were altogether quite rare in AMs (1.6+0.9 = 2.5%), but more abundant in IMs (4.6+5.8 = 10.4%). The final two classes, Class IV and Class V cells were rare and found exclusively in infected IMs. Class IV cells (2.8% of IMs) exclusively expressed viral gRNA, but instead of the discrete puncta (∼1 um in dimension) observed in Classes I – III, formed dense bodies in large regions of the cell. Furthermore, nuclear staining often showed blebbing morphologies, and lysosomal staining was compacted around the nucleus instead of the smooth perinuclear ring seen in the other classes. Class V cells (2.4% of IMs) demonstrated abundant but diffuse viral gRNA expression, and often were lysed or lacked a clearly definable nuclear or lysosomal architecture. These are inferred to be the final stage in IM destruction.

#### Super-resolution (SR) microscopy

For SR imaging, samples were mounted in a blinking-inducing buffer that consists of 200 U/ml glucose oxidase, 1000 U/ml catalase, 10% w/v glucose, 200 mM Tris-HCl pH 8.0, 10 mM NaCl and 50 mM cysteamine. The buffer was prepared from following stock solutions^72^: 1) 4 kU/ml glucose oxidase (G2133, Sigma-Aldrich), 20 kU/ml catalase (C1345, Sigma), 25 mM KCl, 4 mM TCEP, 50% v/v glycerol and 22 mM Tris-HCl pH 7.0, stored at −20 °C; 2) 1 M cysteamine-HCl (30080, Sigma-Aldrich), stored at −20 °C; 3) 40% w/v glucose, stored at RT; 4) 1M Tris-HCl pH 8.0 (J22638.AE, Thermo Fisher Scientific).

SR imaging was performed on a custom inverted epifluorescence microscope as described before^63^ (Nikon Diaphot 200 frame) equipped with an Electron Multiplying Charge-Coupled Device (EMCCD) camera (Andor iXon DU-897) and a 60x/1.4 NA oil-immersion objective (Olympus PLAPON60XOSC2). We used a custom tube lens (f=400mm) that provides a calibrated pixel size of 117.2 nm. Fluorophore emission was collected through a dichroic mirror (Chroma, ZT440/514/561/640rpc-UF1) and filtered by a notch and a bandpass filters (Chroma, ZET642NF and ET700/75m). Alexa Fluor 647 was excited with a 642 nm continuous-wave laser (MPB Communications Inc.) at ∼ 5 kW/cm^2^. For each field, ∼45000 frames were acquired with an exposure time of 20 ms and an EM gain of 100. The single-molecule localizations were detected and fitted to an integrated Gaussian PSF model with a weighted least squares method and corrected for drift by cross-correlation in ThunderStorm^73^. Final SR images were reconstructed as 2D histograms of localizations with a bin size of 20×20nm^2^, using only localizations with fitted sigma between 100 and 180 nm and uncertainties under 30 nm as determined by ThunderStorm.

#### Pseudotyped lentivirus production

To create lenti-S-NLuc-tdT, Spike pseudotyped lentiviruses encoding a nanoluciferase-tdTomato reporter were produced in HEK-293T cells (5 × 10^6^ cells per 10-cm culture dish) by co-transfection of a 5-plasmid system as described previously^74^. Based on the original lentiviral backbone plasmid (pHAGE-Luc2-IRES-ZsGreen, Addgene 164432), we replaced the Luc2-IRES-ZsGreen reporter with a cassette encoding H2B fused to Nanoluciferase (Promega) to minimize background luminescence, followed by a T2A self-cleaving peptide, and tdTomato fluorescent protein using in-Fusion cloning (Takara Bio). The 5-plasmid system includes a packaging vector (pHAGE-H2B-NanoLuc-T2A-tdTomato), a plasmid encoding full-length Spike with a 21-residue deletion on the C-terminus (pHDM SARS-CoV-2-SpikeΔ21), and three helper plasmids (pHDM-Hgpm2, pHDM-Tat1b, and pRC-CMV_Rev1b). Transfection mixture was prepared by adding 5 plasmids (10 µg packaging vector, 3.4 µg Spike-encoding plasmid, and 2.2 µg of each helper plasmid) to 1 mL D10 medium (DMEM supplemented with 10% FBS, 1% Pen/Strep/L-Glutamine), followed by the addition of 30 µL BioT transfection reagent (Bioland Scientific, LLC, B01-03) in a dropwise manner with vigorous mixing. After 10-min incubation at room temperature, the transfection mixture was transferred to HEK-293T cells in the culture dish. Culture medium was replenished 24 hr post-transfection, and after another 48 hr, viruses were harvested and filtered through a 0.45-µm membrane. Spike-pseudotyped lentiviruses were aliquoted, stored at -80°C, and titrated in HeLa/ACE2/TMPRSS2 cells before being used in neutralization assays. For non-pseudotyped control lentivirus (Spike-), transfection was performed similarly except omitting pHDM SARS-CoV-2-SpikeΔ21.

#### Neutralizing antibodies

The variable heavy chain (HC) and variable light chain (LC) sequences for COVA2-15^75^ (HC GenBank MT599861, LC GenBank MT599945) were codon optimized for mammalian expression. Fragments were PCR amplified and inserted into linearized CMV/R expression vectors containing the heavy chain or light chain constant domains from VRC01. COVA2-15 was expressed in Expi293F cells via transient transfection and purified on a MabSelect PrismA column (Cytiva) on an ÄKTA Protein Purification System. Fractions were concentrated, buffer-exchanged to HEPES buffer saline (20 mM, pH 7.4, 150 mM NaCl) with 10% glycerol, flash-frozen in liquid nitrogen and stored at -20°C. REGN antibodies were a gift from D. Burton.

#### Entry and neutralization assays

Lenti-S-NLuc-tdT (Spike+ (D614G)) or a non-pseudotyped control lentivirus (Spike-) (diluted in DMEM/F12 medium, supplemented with polybrene, 1:1000, *v/v*) were added to purified AMs or IMs in culture, and, after 4 hours, free virions were washed off and infections continued for 48 hours before quantification of infection by expression level (luminescence) of the NLuc reporter. Uninfected AMs or IMs (Cells only) served as background control.

Neutralization was performed with the monoclonal antibodies (mAbs) against SARS-CoV-2 Spike described above by pre-incubating them with the lentivirus, or with blocking antibodies against the following cellular receptors: ACE2 (R&D AF933), CD169/Siglec-1 (Clone 7-239, Biolegend 346002), or CD209/DC-SIGN (Clone 120507, R&D MAB161-500). The effect of the antibodies on viral entry into purified macrophages was measured with Spike-pseudotyped lentiviruses with the nanoluciferase-tdTomato reporter. Purified AMs or IMs were seeded in white-walled, clear-bottom 96-well plates (10,000-20,000 cells per well) 1-day before the assay (day 0). On day 1, Spike-targeting mAbs were serially diluted in DMEM/F12 medium and then mixed with lentivirus (diluted in DMEM/F12 medium, supplemented with polybrene, 1:1000, *v/v*) for 1 hr before being transferred to macrophages. Alternatively, macrophages were incubated with blocking antibodies against the cellular receptors, along with FcR blockade (Biolegend Human TruStain FcX, 422302) to prevent FcR-mediated antibody-dependent phagocytosis, for 1 hr before the addition of lentiviruses. Culture medium was replenished 4 hr post-infection. On day 3, medium was removed and cells were rinsed with Dulbecco’s Phosphate-Buffered Saline (DPBS, Gibco, 14190144) before 100 µL nanoluciferase substrate (Nano-Glo, Promega, N1110) were added to each well. Luminescence was recorded on a BioTek Synergy™ HT or Tecan M200 microplate reader. Percent infection was normalized to cells only (0% infection) and virus only (100% infection) on each plate. Neutralization assays were performed in biological duplicates (macrophage purifications from distinct donors).

#### Statistics and Reproducibility

UMAP plots include every cell from indicated cell types, taken from one representative case without performing data integration. All heat maps and plots with single cell expression data include every cell from indicated types, unless otherwise stated in the figure legend (numbers available in Supplementary Table 4). Dot plots were generated using a modified version of Scanpy’s ‘pl.dotplot’ (Fig. 2a, viral expression dot plot) with indicated expression cutoff, or a modified version of Seurat’s ‘DotPlot’ function available on GitHub (in Figs. 3d, 4g, and Extended Data Fig. 6). Scatter plots for infection pseudotime were generated with ggplot2’s ‘geom_point’ function, and trend lines were plotted with parameter ‘method = “loess”’ (Figs. 3g, 4da-f, Extended Data Fig. 5e). Violin plots were generated with Scanpy’s ‘pl.violin’ function (Fig. 1c, left panel), or Seaborn’s ‘violinplot’ and ‘stripplot’ functions (Fig. 2a) and show proportion of single cells at indicated expression levels. Bar plots were generated in Excel (Fig. 1c, right panel, 3f). Histogram plots were generated using Seaborn’s ‘histplot’ function with log scale transformation on both x-axis and y-axis (Fig. 1d, lower panel). Cumulative distribution plot was generated using Seaborn’s ‘ecdfplot’ function and plotted on a Matplotlib’‘logit’ scale which implements the logistic distribution (in Fig. 1d, upper panel). Arcplots depicting number of subgenomic junctions was plotted using a custom Python function (available on Github). Differentially expressed genes along infection pseudotime were computed by taking the top 250 genes that contributed to each pseudotime trajectory (see Methods), and further tested using pseudotimeDE’s ‘runPseudotimeDE’ function without subsampling testing against the asymptotic null distribution, with exact p-values indicated in Table S3. Differentially expressed genes between “Late” vs. “Early” macrophages along infection pseudotime were computed using Seurat’s ‘FindMarkers’ function implemented using the default Wilcoxon Rank Sum test, with exact p-values indicated in Table S3. Immunostaining and smFISH experiments were performed on at least 2 human or mouse subjects distinct from the donors used for sequencing, and quantifications were based on at least 10 fields of view in each. For smFISH, fields of view were scored manually, calling a cell positive for each gene probed if its nucleus had at least three associated expression puncta. No statistical methods were used to predetermine sample size. The experiments were not randomized and investigators were not blinded to allocation during experiments and outcome assessment.

### Data Availability

Raw sequencing data, UMI tables, cellular annotation metadata, Seurat objects, and scanpy objects are being deposited and will be released as soon as possible (at latest, upon acceptance of this manuscript).

### Code Availability

Code to reproduce the analyses and figures are being deposited and will be released as soon as possible (at latest, upon acceptance of this manuscript).

## Acknowledgements

We are grateful to the lung donors and Donor Network West for whole lung tissue, and to an anonymous financial donor for construction of a new BSL3 facility. We thank Amy Kistler (Chan-Zuckerberg Biohub) for viral stock sequencing assistance; Jaishree Garhyan, Geoffrey Ivison, and Michelle Leong (Stanford) for BSL3 support; Daphne Cooper (10x Genomics) for discussions on BSL3 implementation of 10x protocols; and Robert C. Jones (Stanford) for help in tissue and equipment procurement. We also thank members of the Blish and Krasnow labs for valuable discussions and comments on the manuscript, and Alexander Lozano for discussions on bioinformatic analyses. The SARS-Related Coronavirus 2, Isolate USA-WA1/2020, NR-52281 was deposited by the Centers for Disease Control and Prevention and obtained through BEI Resources, NIAID, NIH. Funding was provided by the Bill & Melinda Gates Foundation (C.A.B., M.A.K.), Chan Zuckerberg Biohub (C.A.B., S.R.Q., P.S.K.), the Burroughs Wellcome Fund Project 1016687 (C.A.B.), a Stanford Chem-H/Innovative Medicine Accelerator COVID-19 Response Award (C.A.B.), and the Howard Hughes Medical Institute (M.A.K.). Fellowship and training support was from National Institutes of Health (A.R., T32 AI007502 and K08 AI163369; G.M-C., T32 DK007217; A.W., T32 GM007364), American Cancer Society Postdoctoral Fellowship (S.K.J.), Stanford Maternal & Child Health Research Institute (D.X.), Stanford Graduate Fellowship and Stanford Cell and Molecular Biology Training Grant (T32GM007276) (Y.Z.), and Stanford Bio-X Interdisciplinary Graduate Fellowship (T.H.W., A.W.). D.D.L. and A.W. were supported by Stanford University Medical Scientist Training Program grant T32-GM007365 and T32-GM145402. C.A.B. is an investigator of the Chan Zuckerberg Biohub and M.A.K. is an investigator of the Howard Hughes Medical Institute.

## Author Contributions

T.H.W., K.J.T, A.R., A.G., S.Q., C.K., C.A.B., and M.A.K. conceived the project and designed the lung slice culture, SARS-CoV-2 infection, lung cell isolation, scRNA-seq, and bioinformatic analysis strategy. T.H.W., W.T., J.S. and C.S.K. reviewed clinical or donor histories and coordinated patient care teams to obtain profiled tissues. J.B.S. and C.S.K. established and revised IRB protocol 15166 to optimize for single cell and lung slice culture preparations. A.R., G.M-C,, and C.A.B. built the BSL3 SARS-CoV-2 program at Stanford University. A.R., G.M.-C. and A.B. isolated, propagated, genotyped, titered, sequenced, and developed inactivation and inhibition methods for SARS-CoV-2 virions and fixation methods for infected tissues. A.R. performed SARS-CoV-2 infections, and plaque assays. T.H.W., K.J.T, A.R., Y.Z., S.K.J., A.G., G.M-C, A.B., A.J.W., and M.M. designed and processed tissue to single cell suspensions, and prepared sequencing libraries. T.H.W. and K.J.T processed and aligned sequencing data. T.H.W., K.J.T, A.R., Y.Z., S.K.J., and A.G. provided tissue expertise and annotated cell types. R.D. performed SARS-CoV-2 sequence alignment, implemented SICILIAN, and with T.H.W., K.J.T., and R.B. analyzed subgenomic junction results. T.H.W. and K.J.T implemented bioinformatic methods, including infection pseudotime. T.H.W., K.J.T and S.K.J. designed, performed, and analyzed multiplex in situ hybridization experiments. T.H.W. and D.D.L developed and implemented the macrophage purification strategy. T.H.W. performed HCR in situ hybridization and immunofluorescence staining of infected macrophages, and L.A. performed super-resolution microscopy. T.H.W., A.R., and D.X. designed, performed, and analyzed Spike-dependent macrophage entry experiments. I.L.W., J.B.S., S.R.Q., C.S.K., J.S., W.E.M., P.S.K., A.R., C.A.B. and M.A.K. supervised and supported the work. T.H.W., K.J.T, A.R., C.A.B., and M.A.K. interpreted the data and wrote the manuscript, and all authors reviewed and edited the manuscript.

## Extended Data Figure Legends

**Extended Data Figure 1.**
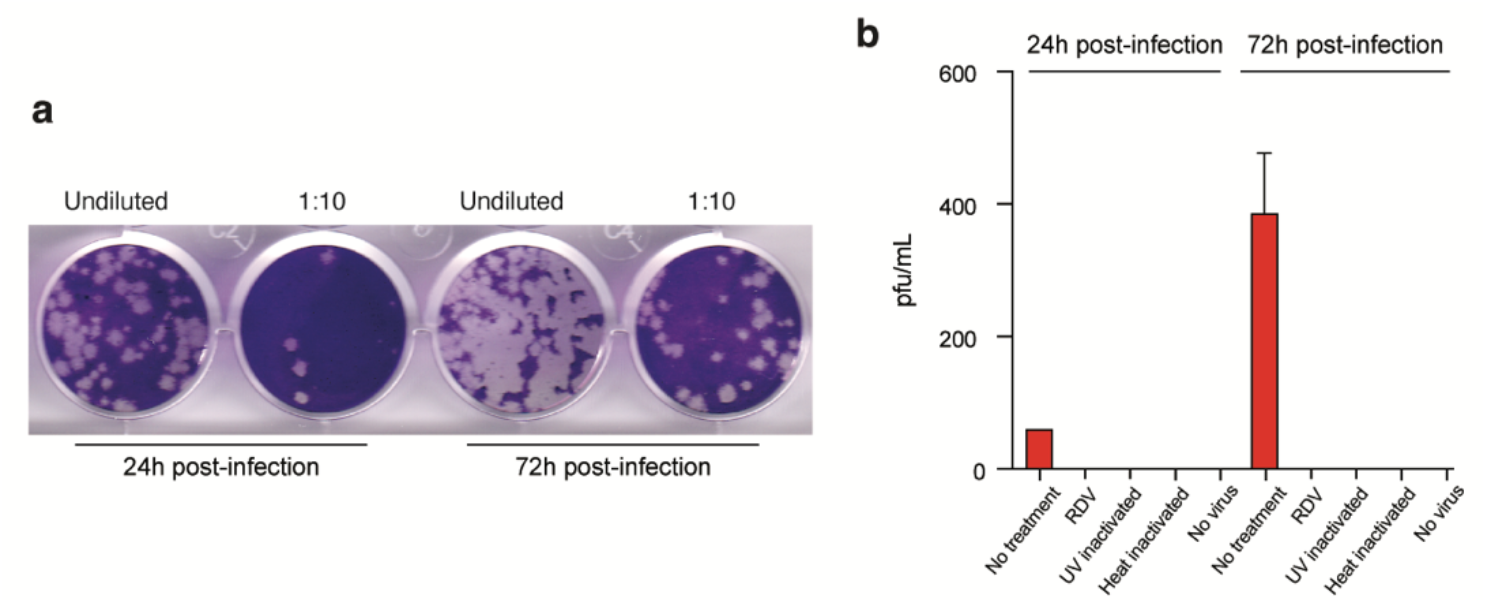
Virion production of cultured human lung slices infected by SARS-CoV-2 increases over time. (a) Plaque assays on VeroE6/TMPRSS2 cells as in Fig. 1 of supernatant collected serially at 24 hr and 72 hr from the same lung slice culture. Media was completely replaced after the harvest of supernatant at 24 hr. (b) Quantification of (a) showing plaques at 24 and 72 hr, along with similar quantification of plaque assay results for remdesivir (RDV) treatment, UV viral inactivation, heat inactivation, and no virus controls (as in Fig. 1). Values shown are the mean+SD of technical duplicates of the plaque assay. pfu, plaque forming units.

**Extended Data Figure 2.**
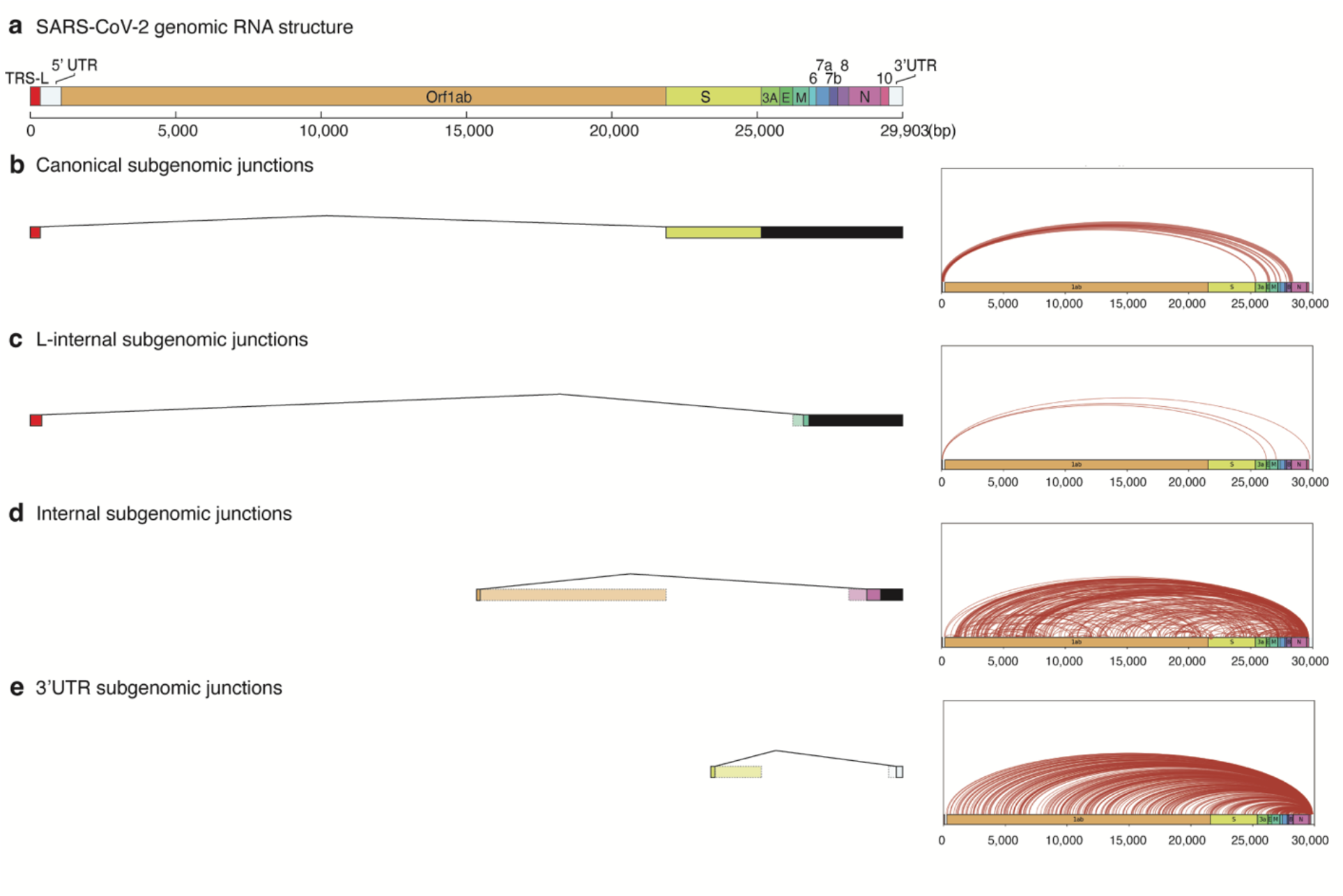
Classes and abundance of canonical and novel subgenomic junctions detected in cultured human lung slices infected by SARS-CoV-2. SARS-CoV-2 subgenomic RNA junctions were identified in scRNAseq analysis of infected lung slice cultures from lung slices infected in all cases as individual sequence reads that mapped discontinuously on the viral genome, as called by SICILIAN (SIngle Cell precIse spLice estImAtioN)^43^ using generalized linear statistical modeling for precise unannotated splice junction quantification in single cells. (a) Diagram of full-length SARS-CoV-2 genomic RNA (29,903 nt) showing annotated ORF positions, the common 5’ “leader” transcription-regulatory sequence (TRS-L, red fill) that connects in viral subgenomic RNAs to gene body TRS-B elements (not shown) adjacent to each of the canonically recognized ORFs (other colors), and the 5’ and 3’ untranslated regions (UTRs, open fill) of the viral genome. (b-d) Examples of inferred subgenomic RNA structures (left panel) based on the type of subgenomic junction detected, alongside arc plots (right panel) visualizing all novel junctions detected for that subgenomic junction type across all infection replicates. (b) “Canonical” subgenomic junctions connect the common 5’ “leader” transcription-regulatory sequence (TRS-L) to gene body (TRS-B) adjacent to each of the canonically recognized ORFs. (c-e) “Noncanonical” subgenomic junctions, which are consistent with previous long read sequencing results from in vitro infections of diverse cell lines by different viral isolates^71,76^. (c) Rare “L-internal” junctions connect TRS-L to cryptic gene body fusion sites. These could represent aberrant jumps during discontinuous transcription. (d) “Internal” junctions occur between any two internal sites within the gene body. (e) The most abundant “3’UTR” junctions occur between any internal site within the gene body and the 3’ UTR of the genome. These are likely overrepresented due to the predominant bias in sequence reads to the 3’-end in the scRNAseq technology employed (10x Genomics).

**Extended Data Figure 3.**
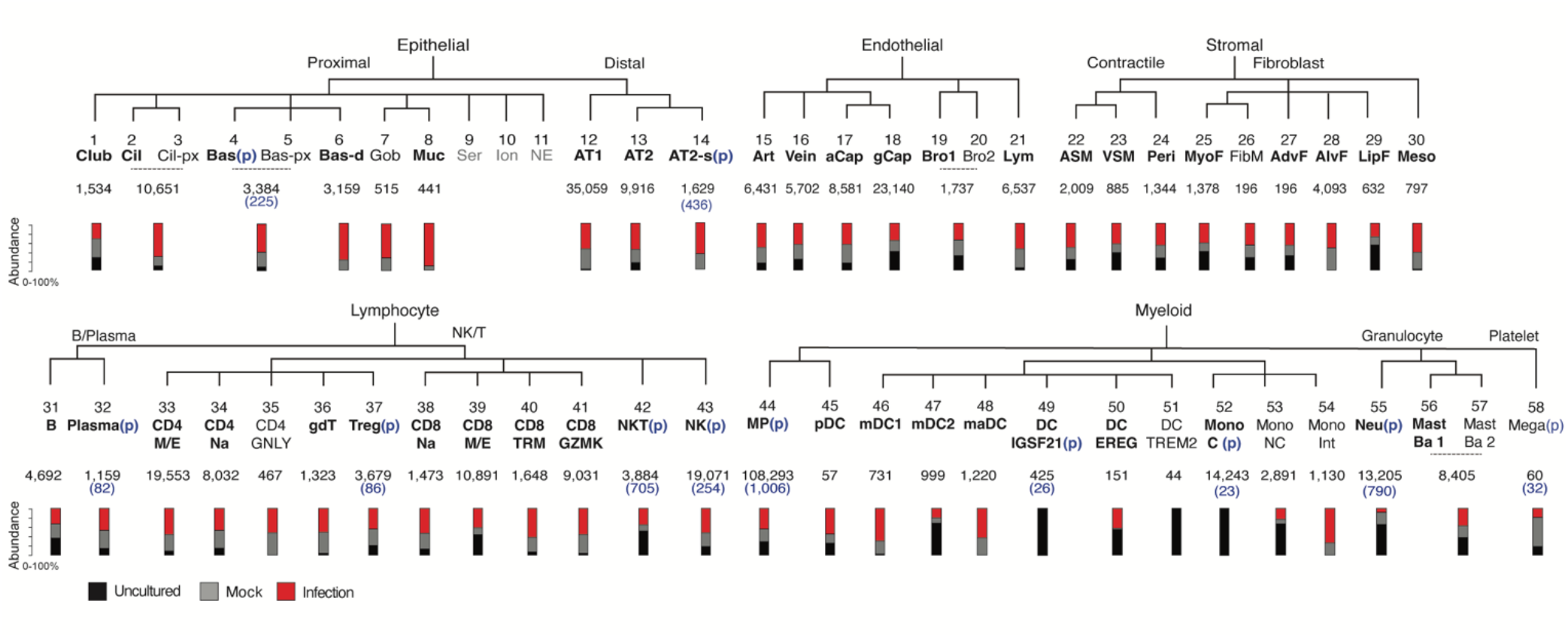
Identity and abundance of canonical and novel lung cell types detected in human lung slice cultures by scRNA-seq. Hierarchical tree showing human lung molecular cell types and their annotations in the indicated tissue compartments following iterative clustering of scRNA-seq profiles of cells from cases 1-4 in each compartment. Numbers below cell type name show total abundance of the cell type, and the stacked bar plot indicates proportions detected from each condition of freshly profiled uncultured (Uncultured), cultured and mock infected (Mock), or cultured and infected (Infected). Black, canonical cell types per our healthy reference human lung cell atlas^34^ (bolded, detected in > 1 lung slice dataset). Cell types in which a proliferative subpopulation was detected is indicated (p) with the number of proliferative cells given in parenthesis. Cell types that were difficult to distinguish via 10x expression profiles without full-length transcriptome were merged. Abbreviations: Cil, ciliated; Cil-px, proximal ciliated; Bas, basal; Bas-px, proximal basal; Bas-d, differentiating basal; Gob, goblet; Ser, serous; Ion, ionocyte; NE, neuroendocrine; AT1, alveolar epithelial type 1; AT2, alveolar epithelial type 2; AT2-s, signaling alveolar epithelial type 2. Art, artery; aCap, capillary aerocyte; gCap, general capillary; Bro, bronchial vessel; Lym, lymphatic. ASM, airway smooth muscle; VSM, vascular smooth muscle; Peri, pericyte; MyoF, myofibroblast; FibM, fibromyocte; AdvF, adventitial fibroblast; AlvF, alveolar fibroblast; LipF, lipofibroblast; Meso, mesothelial. CD4 M/E, CD4 memory/effector T cells; CD4 Na, CD4 naïve T cells; Treg, regulatory T cells; CD8 TRM, CD8 tissue resident memory T cells; NK, natural killer cell. MP, macrophage; pDC, plasmacytoid dendritic cell; mDC, myeloid dendritic cell; maDC, mature dendritic cell; Mono C, classical monocyte; Mono NC, nonclassical monocyte; Mono Int, intermediate monocyte; Neu, neutrophil; Mast Ba, mast/basophil; Mega, megakaryocyte.

**Extended Data Figure 4.**
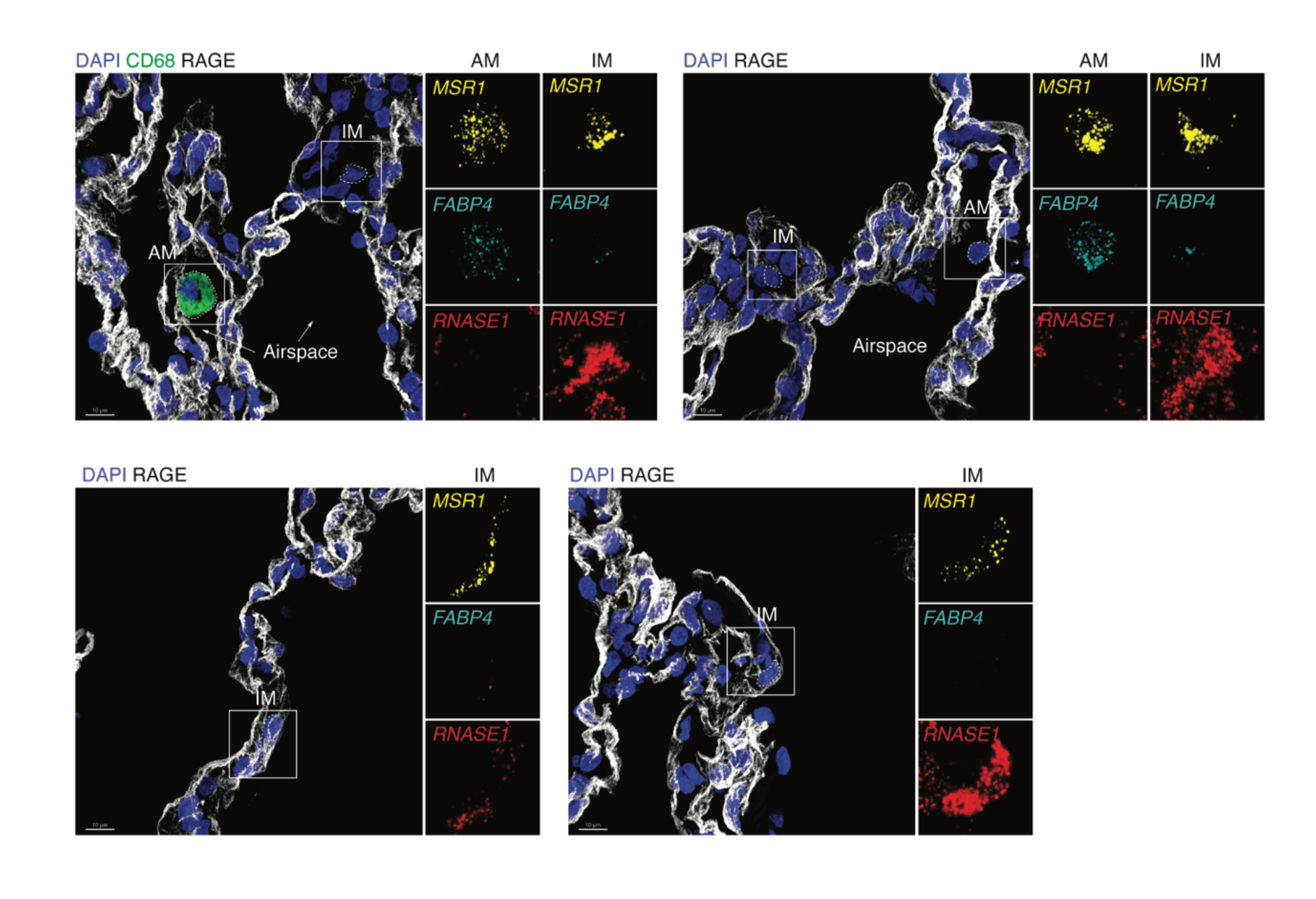
Localization and morphology of interstitial and alveolar macrophages in the lung. Additional examples as in Fig. 3e of RNAscope single molecule fluorescence in situ hybridization (smFISH) and immunostaining of alveolar (AM) and interstitial (IM) macrophages in non-cultured human lung of case 2, detecting general macrophage antigen CD68 (green, protein), AT1 antigen RAGE (white, protein), AM marker *FABP4* (cyan, RNA), and IM marker *RNASE1* (red, RNA). Scale bar, 10 µm.

**Extended Data Figure 5.**
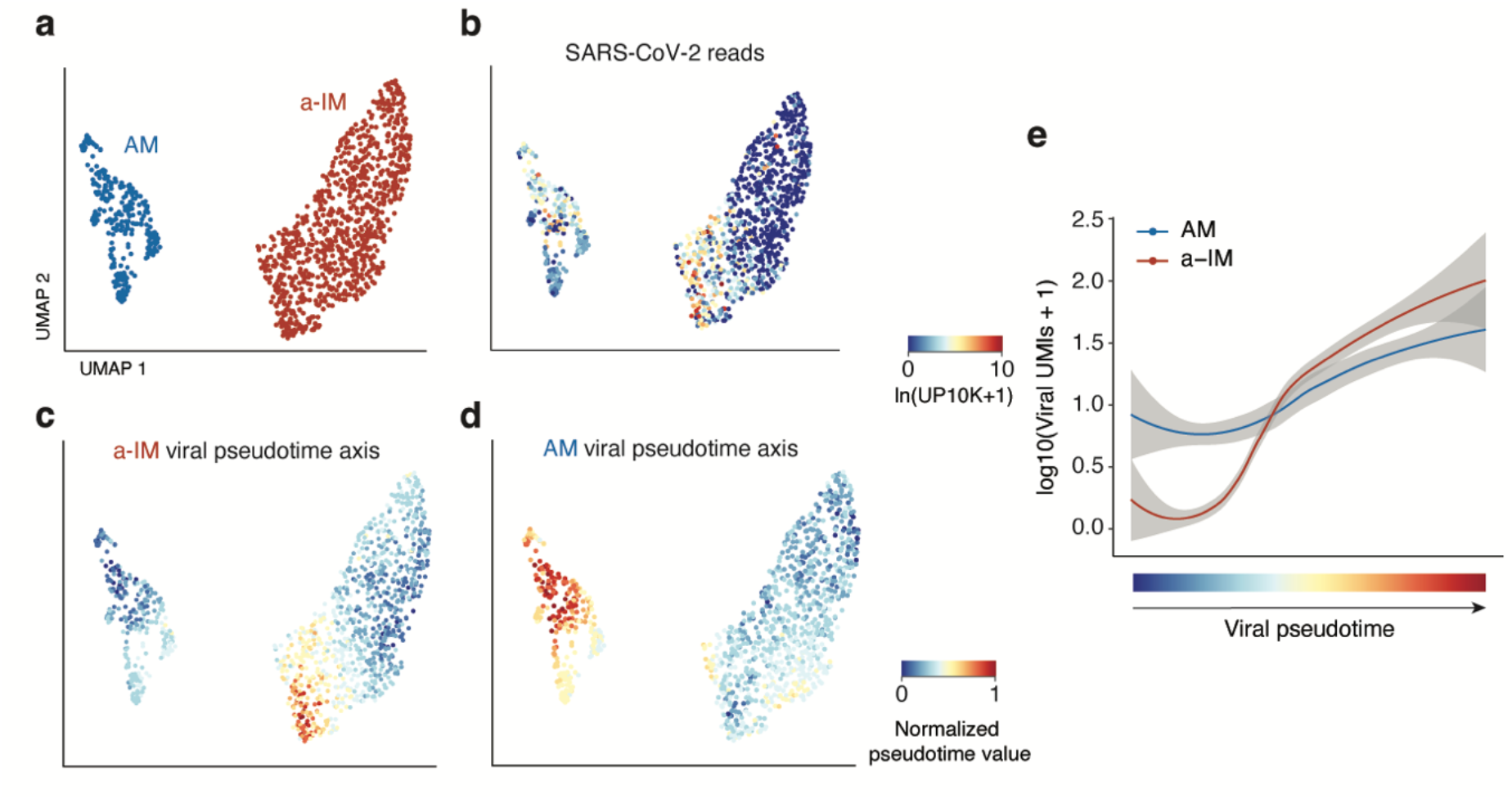
Distinct viral pseudotime trajectories in interstitial and alveolar macrophages. (a) Uniform Manifold Approximation and Projection (UMAP) projection of alveolar macrophages (AM) and activated interstitial macrophages (a-IM) in infected human lung slices from 10x scRNA-seq from infections 1 and 4, as in Fig. 3. (b) Normalized expression of SARS-CoV-2 RNA in each cell as shown by the heat map scale. (c) Normalized value of a-IM viral pseudotime value as shown in Fig. 4a. (d) Normalized value of AM viral pseudotime value as shown in Fig. 4a. (e) Total viral RNA expression (log10(Viral UMIs + 1)) graphed against viral infection pseudotime for AMs and a-IMs.

**Extended Data Figure 6.**
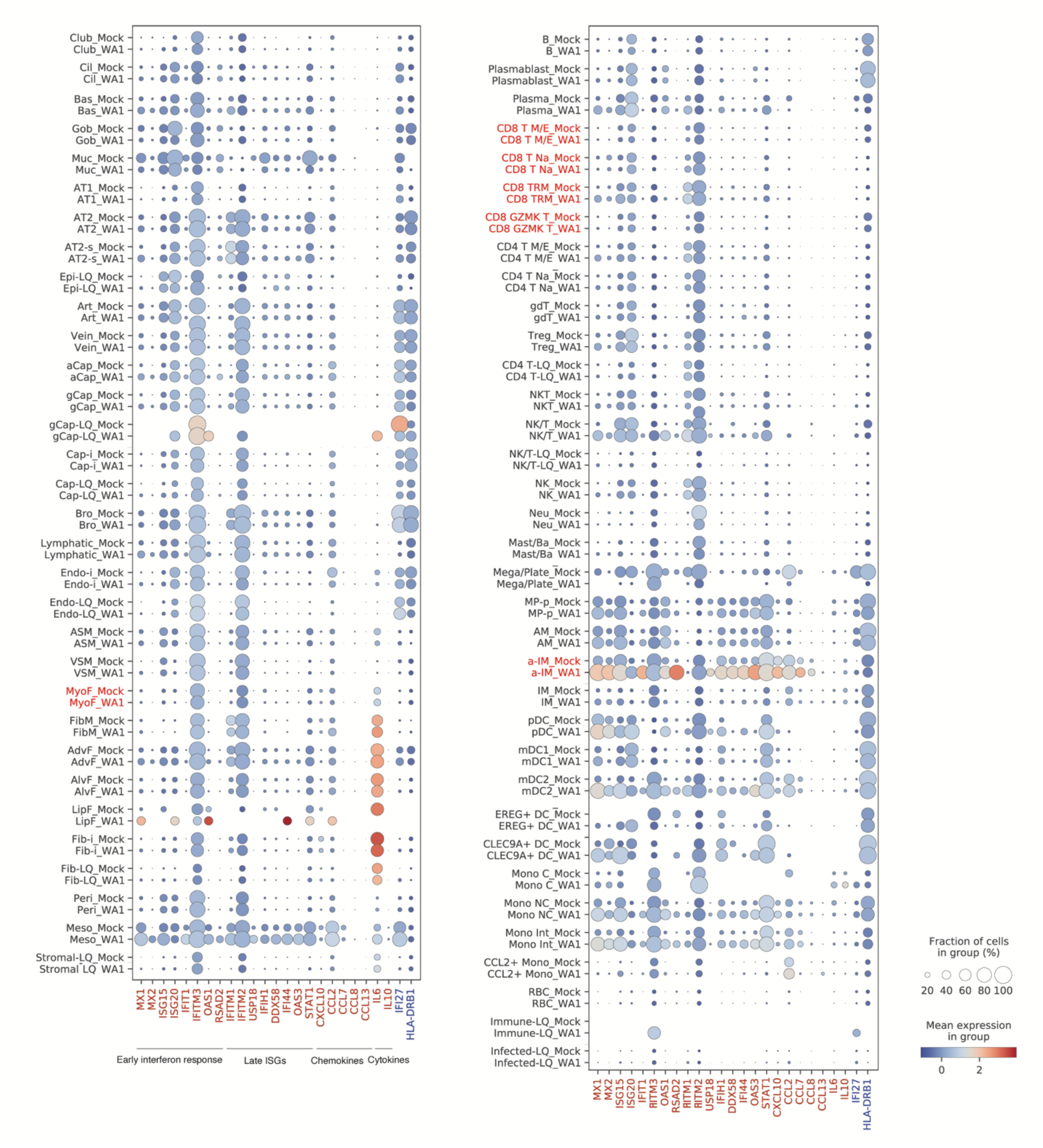
The organ-wide landscape of the a-IM anti-viral inflammatory response in infected human lung slice cultures. Dot plot visualizing the expression of genes (including early interferon response and late interferon stimulated genes (ISGs), chemokine and cytokine ligands) that had statistically significant association with infection pseudotime in a-IM (red text) or AMs (blue text) as in Fig. 4, in each of the molecular cell types detected by scRNA-seq of cultured (“_Mock”) or infected (“_WA1”) human lung slice cultures from cases 1 and 4. Note the low but appreciable baseline expression of the antiviral response (early interferon response and late ISGs) in diverse cell types even without viral infection, but the specific induction of it in a-IMs in infected vs. mock cultures. Chemokine and cytokines are expressed at low levels and are better appreciated at single cell resolution (see Extended Data Fig. 7).

**Extended Data Figure 7.**
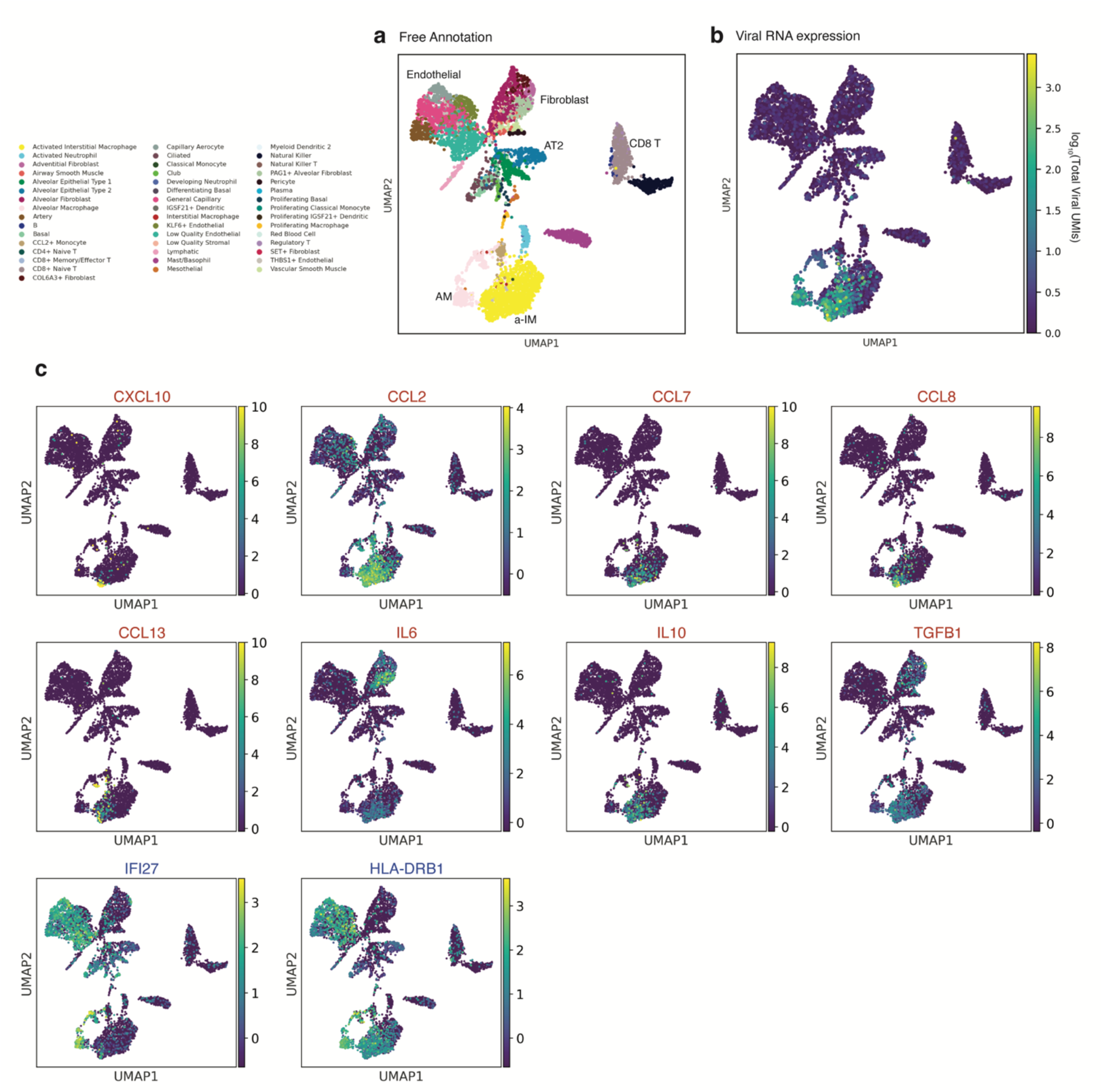
The organ-wide landscape of the a-IM inflammatory signals in infected human lung slice cultures. UMAP visualizing the molecular cell types, SARS-CoV-2 viral load, and expression of the inflammatory chemokine and cytokine ligands that had statistically significant association with infection pseudotime in a-IM (red text), or AMs (blue text) as in Fig. 4, in each of the molecular cell types detected by scRNA-seq of infected human lung slice cultures from case 1. Note the focus of chemokine and cytokine expression that colocalizes with the single cells that express high viral RNA levels.

**Extended Data Figure 8.**
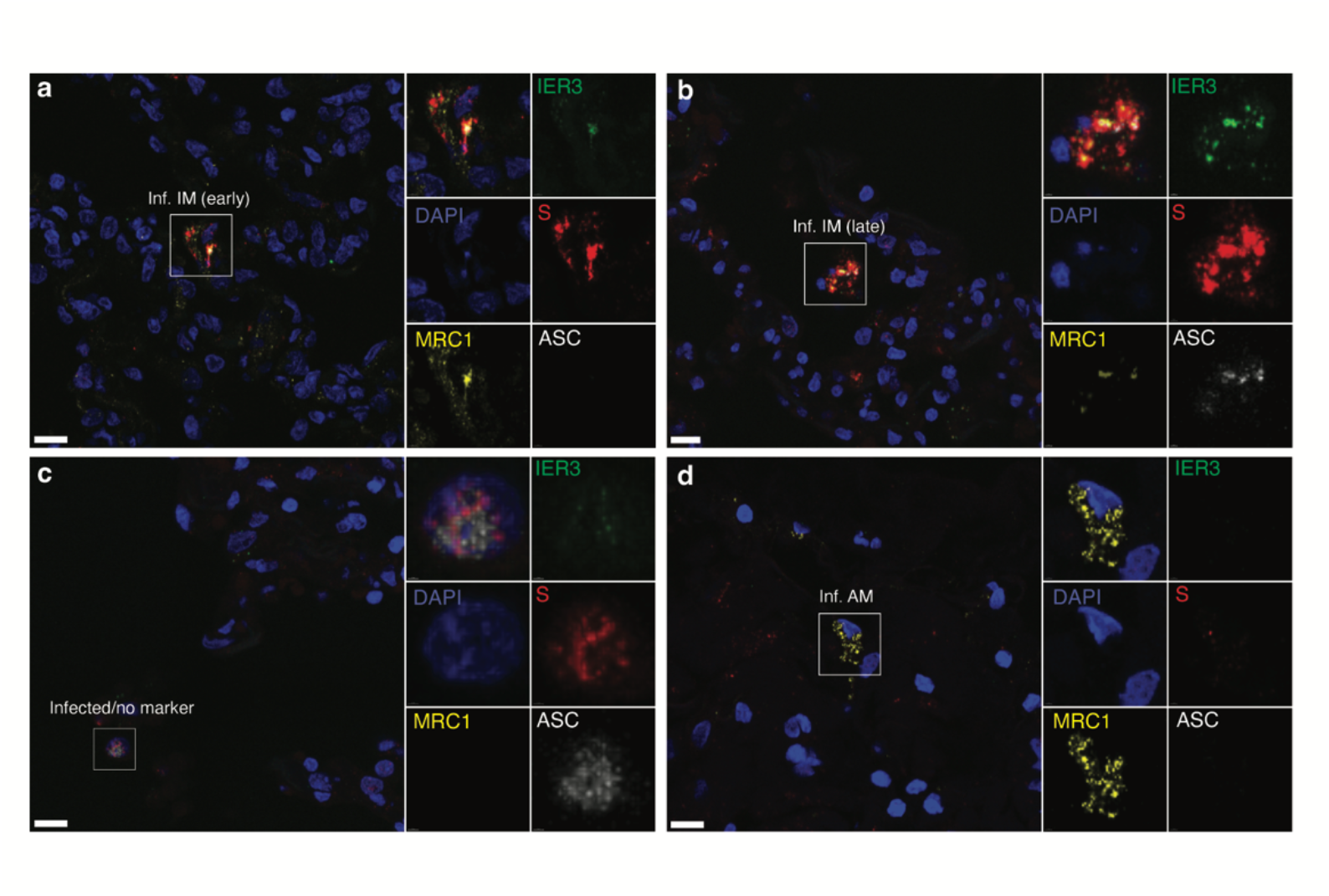
Inflammasome activation in infected human lung slice cultures. Micrographs of RNAscope single molecule fluorescence in situ hybridization (smFISH) and immunostaining of an infected human lung slice culture 72 hrs after infection (as in Fig. 2b) examining inflammasome activation state (ASC speck^+^) of infected alveolar (AM, MRC1^+^IER3^-^ S^+^) and interstitial (IM, MRC1^+^IER3^+^S^+^) macrophages, and terminally infected cells that express none of these markers. ASC immunostaining was rare overall, but could be occasionally detected in late infected IMs (panel b), and late infected “no marker” cells (panel c), marked by abundant Spike staining, and presence of ASC specks.

**Extended Data Figure 9.**
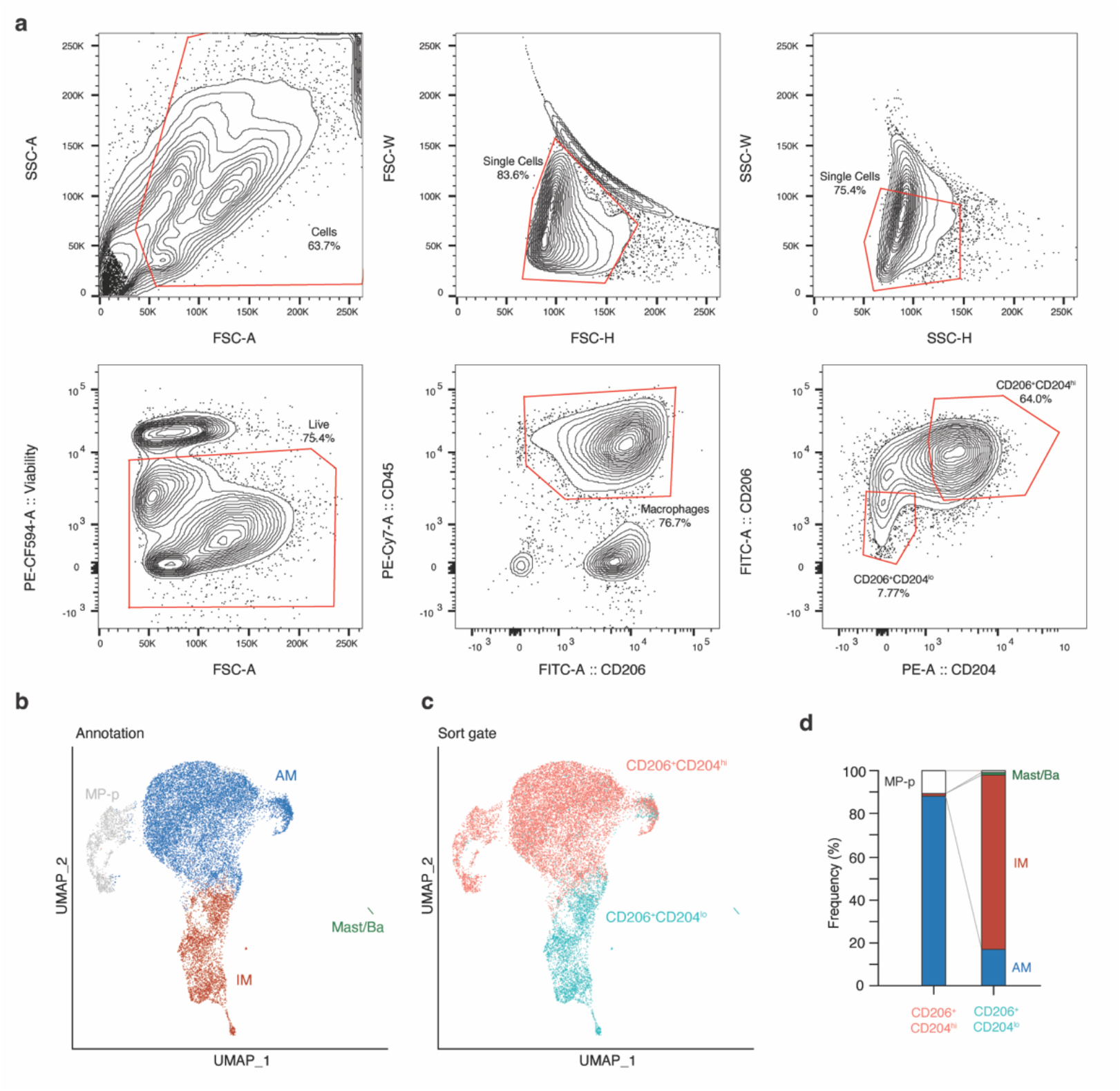
Fluorescent activated cell sorting (FACS) and scRNA-seq of purified AMs and IMs. (a) Sequential FACS data and sorting gates (red) for dissociated human lung cells, following MACS enrichment of lung resident macrophage (MACS CD206^+^) cells. Cells were first gated on viable single cells that were CD45^+^ and CD206^+^ (center bottom panel), then two gates were subsequently sorted (right bottom panel) for 10x scRNA-seq transcriptomic profiling: CD206^+^CD204^hi^ (putative AMs), and CD206^+^CD204^lo^ (putative IMs). (b) Uniform Manifold Approximation and Projection (UMAP) projection of sorted putative AMs and putative IMs from (a), with the transcriptomic molecular cell annotations indicated, including AMs, IMs, proliferating macrophages, and rare Mast/Basophils. (c) The same UMAP projection colored by sorting gate metadata. Note the correspondence between the scRNA-seq molecular annotation and the gating metadata. (d) The relative frequencies of the molecular types of AMs and IMs in each of the indicated sorting gates; in the CD206^+^CD204^hi^ channel, AMs were 88%, IMs were 1%, and proliferating macrophages were 11%; in the CD206^+^CD204^lo^ channel, IMs were 81%, AMs were 17%, proliferating macrophages were 1%, and Mast/Basophils were 1%.

**Extended Data Figure 10.**
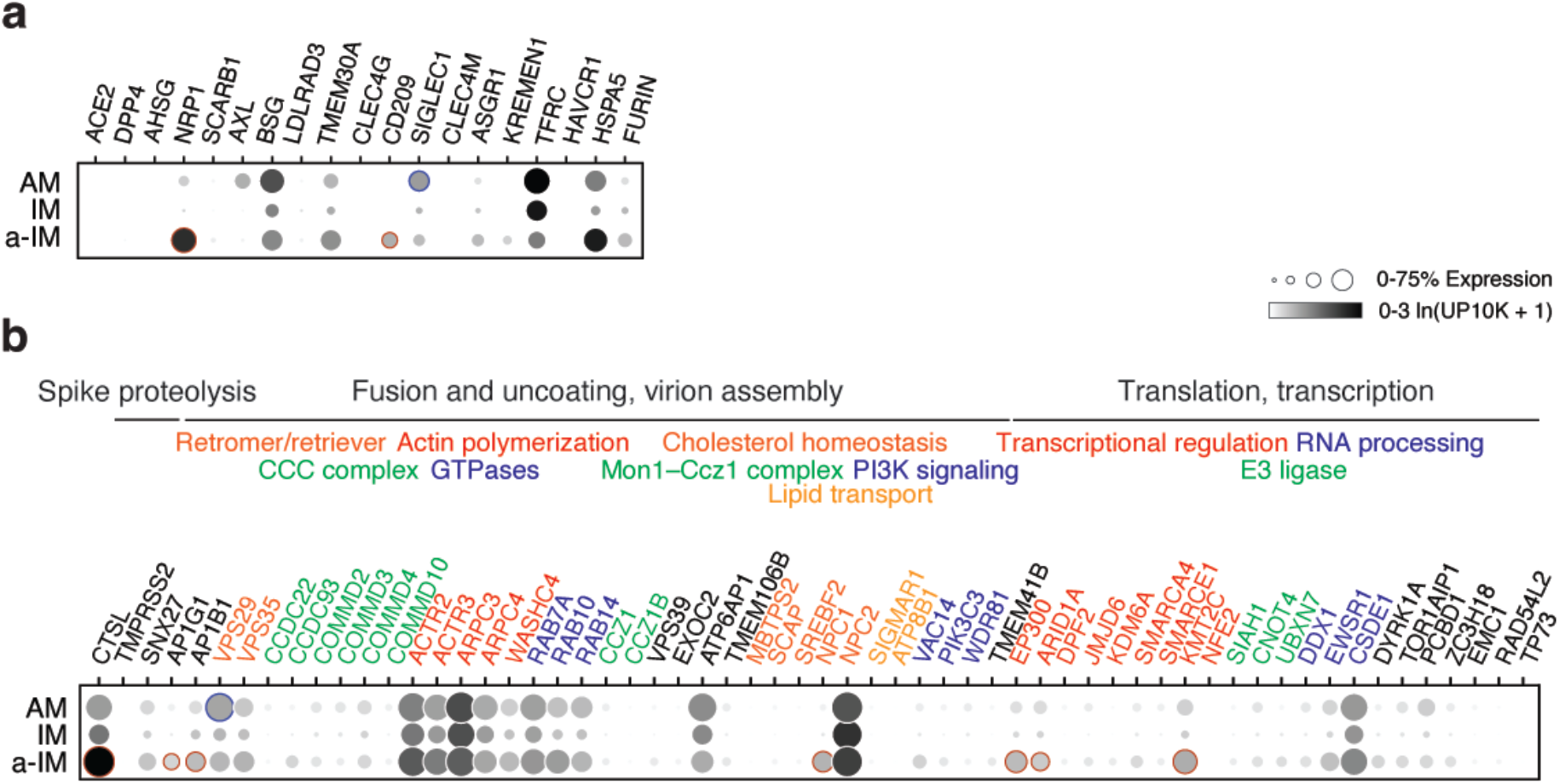
Expression of receptors and other SARS-CoV-2 host factors in different lung macrophage subtypes. Dot plot of scRNA-seq results of freshly profiled human lung slice cultures from cases 1 and 4, as in Fig. 3 showing for each indicated macrophage subtype (AM, alveolar macrophage; IM, interstitial macrophage; a-IM, activated interstitial macrophage) the fraction of expressing cells (% Expression) and mean expression value among expressing cells (ln(UP10K+1) of: (a) proposed canonical and alternative cellular receptors, and (b) other key proviral host factors in the SARS-CoV-2 replication cycle previously identified in CRISPR-based functional genetic screens^65^. Genes are grouped based on different steps of the viral life cycle (black font) and their normal cellular functions (colored font). Dots representing genes differentially up-regulated in a-IMs are outlined in red, and dots representing genes differentially up-regulated in AMs are outlined in blue (adjusted p-value < 0.05). Note differential expression of DC-SIGN/CD209 in a-IMs (including induction relative to IMs), and Sialoadhesin/CD169 in AMs.

**Extended Data Figure 11.**
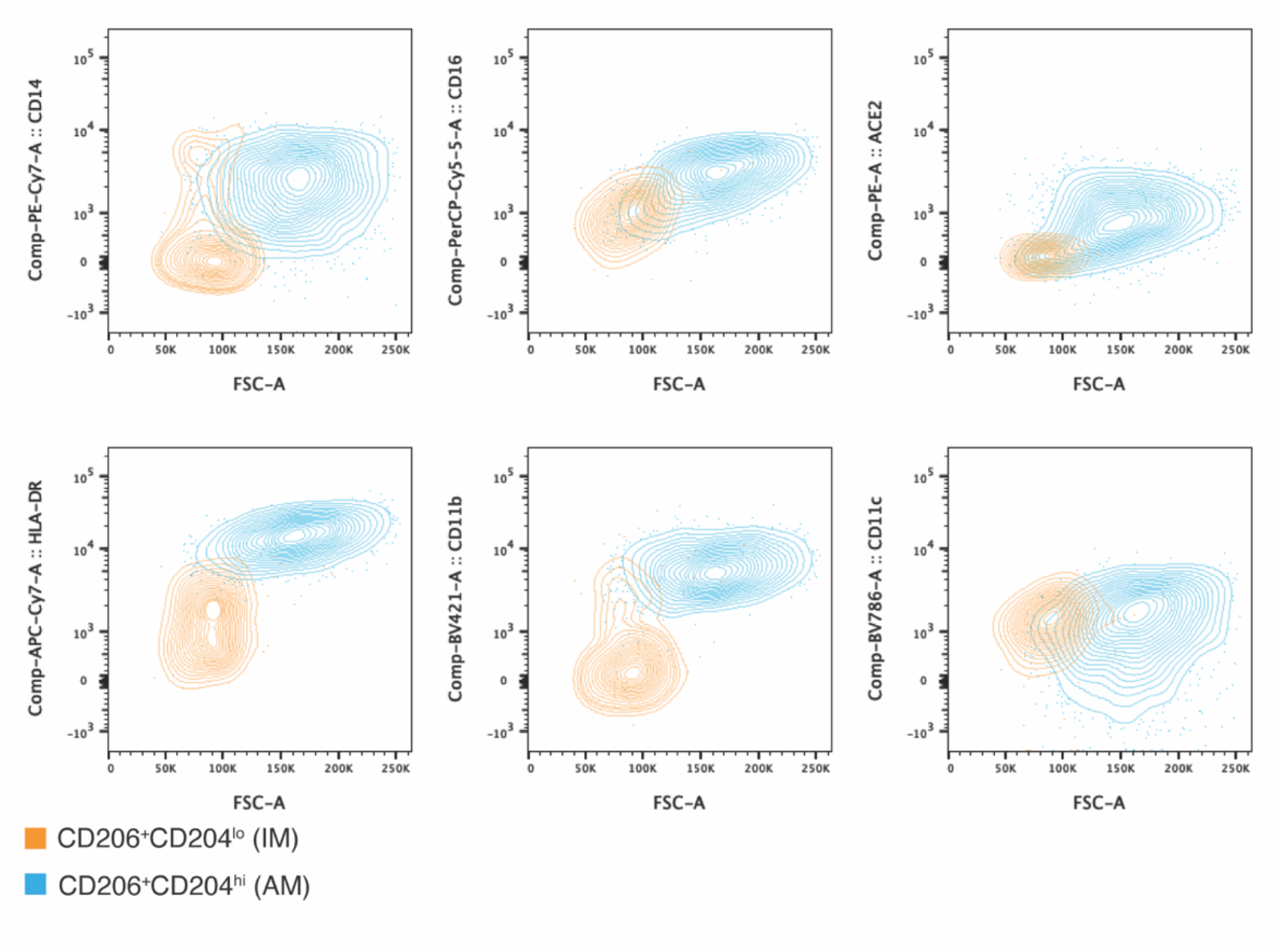
Flow cytometry characterization of AM and IMs collected from freshly dissociated human lung. Flow cytometry of the sorted alveolar macrophages (AMs) and interstitial macrophages (IMs), with flow gating defined as in Extended Data Fig. 9. Results are shown for staining of various surface antigens reported to distinguish AMs and IMs, including CD14 (upper left), CD16 (center top), HLA-DR (left bottom), CD11b (center bottom), and CD11c (lower right), as well as for staining of the canonical SARS-CoV-2 receptor ACE2 (upper right). Note that although neither AMs nor IMs express ACE2 mRNA (Extended Data Fig. 10), AMs, but not IMs, express ACE2 protein.

**Extended Data Figure 12.**
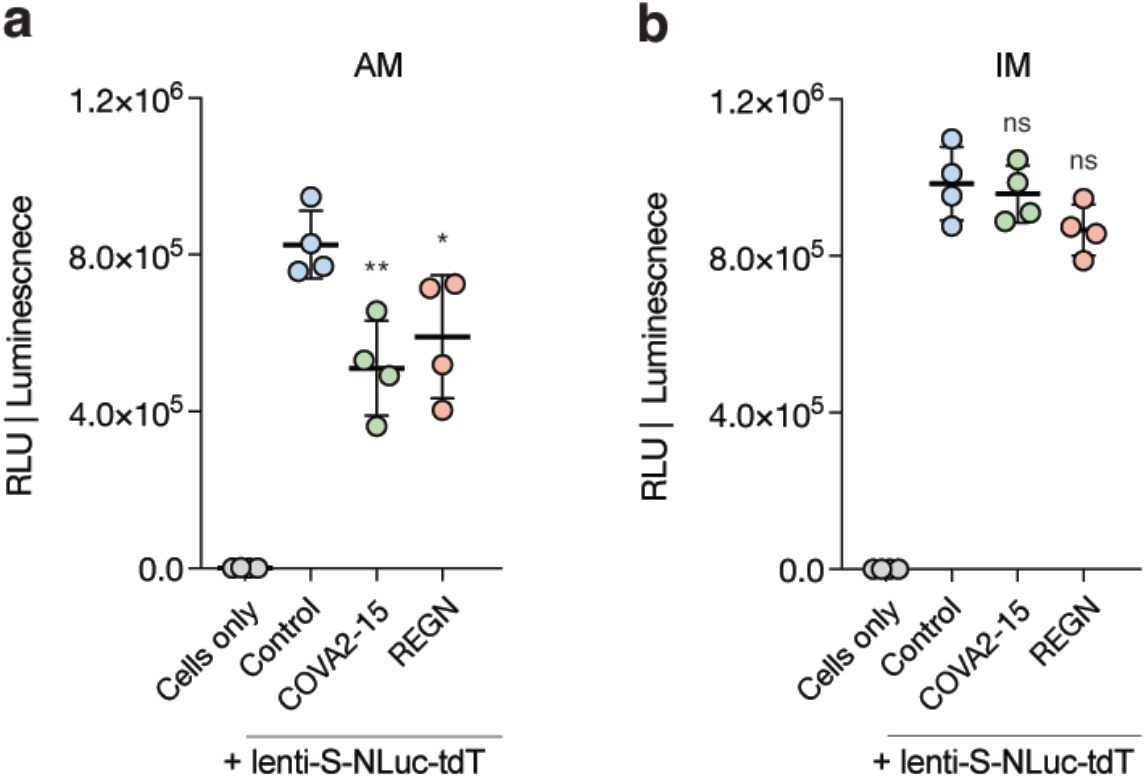
Effect on macrophage entry of therapeutic mAbs targeting the SARS-CoV-2 receptor binding domain. Luminescence readout (RLU, relative light units) of neutralization of SARS-CoV-2 pseudovirus (lenti-S-NLuc-tdT, diluted in DMEM/F12 medium, supplemented with polybrene, 1:1000, *v/v*) by 0.1 ug/ml of the indicated monoclonal antibodies (mAbs) against SARS-CoV-2 receptor binding domain (RBD) in cultured purified alveolar macrophages (AMs) or interstitial macrophages (IMs). To allow mAb binding, virions were pre-treated with the mAb for 1 hr before addition of virions to the cells. NLuc luciferase values are presented as mean ± s.d from two independent experiments. **, P<0.01; *, P<0.05; n.s., non-significant vs. control (no antibody) value.

## Supplemental Tables

**Table S1.** Summary of viral subgenomic junction discovery from single lung cells using SICILIAN.

| GeneA | JuncPosR1A | GeneB | JuncPosR1B | Junction_type | numReads |
| --- | --- | --- | --- | --- | --- |
| SARS-CoV-2_S | 21761 | SARS-CoV-2_S | 21789 | Internal | 168 |
| SARS-CoV-2_L | 64 | SARS-CoV-2_ORF8 | 28255 | Canonical | 120 |
| SARS-CoV-2_N | 29370 | 3'UTR | 29797 | 3'UTR | 82 |
| SARS-CoV-2_ORF6 | 27270 | 3'UTR | 29716 | 3'UTR | 69 |
| SARS-CoV-2_L | 64 | SARS-CoV-2_E | 26468 | Canonical | 69 |
| SARS-CoV-2_ORF6 | 27360 | 3'UTR | 29795 | 3'UTR | 57 |
| unknown | 9462 | 3'UTR | 29706 | 3'UTR | 54 |
| SARS-CoV-2_ORF6 | 27264 | SARS-CoV-2_ORF6 | 27291 | Internal | 47 |
| 3'UTR | 29710 | 3'UTR | 29793 | 3'UTR | 45 |
| 3'UTR | 29685 | 3'UTR | 29804 | 3'UTR | 44 |
| SARS-CoV-2_ORF10 | 29633 | SARS-CoV-2_ORF10 | 29669 | Internal | 42 |
| SARS-CoV-2_S | 25008 | 3'UTR | 29702 | 3'UTR | 40 |
| SARS-CoV-2_N | 28735 | 3'UTR | 29783 | 3'UTR | 33 |
| 3'UTR | 29710 | 3'UTR | 29791 | 3'UTR | 32 |
| 3'UTR | 29711 | 3'UTR | 29777 | 3'UTR | 31 |
| SARS-CoV-2_N | 29361 | 3'UTR | 29801 | 3'UTR | 28 |
| SARS-CoV-2_ORF7b | 27876 | 3'UTR | 29737 | 3'UTR | 28 |
| SARS-CoV-2_L | 66 | SARS-CoV-2_ORF6 | 27385 | Canonical | 27 |
| unknown | 1383 | SARS-CoV-2_N | 29494 | Internal | 27 |
| 3'UTR | 29698 | 3'UTR | 29775 | 3'UTR | 24 |
| SARS-CoV-2_L | 65 | SARS-CoV-2_S | 25381 | Canonical | 24 |
| SARS-CoV-2_L | 69 | SARS-CoV-2_ORF5 | 27041 | Canonical | 24 |
| unknown | 3247 | 3'UTR | 29703 | 3'UTR | 23 |
| unknown | 4647 | 3'UTR | 29782 | 3'UTR | 22 |
| unknown | 4972 | 3'UTR | 29716 | 3'UTR | 22 |
| SARS-CoV-2_ORF5 | 27190 | SARS-CoV-2_ORF6 | 27222 | Internal | 22 |
| SARS-CoV-2_N | 28289 | 3'UTR | 29801 | 3'UTR | 20 |
| unknown | 6247 | 3'UTR | 29767 | 3'UTR | 20 |
| 3'UTR | 29728 | 3'UTR | 29760 | 3'UTR | 19 |
| unknown | 11375 | 3'UTR | 29771 | 3'UTR | 18 |
| unknown | 15193 | 3'UTR | 29709 | 3'UTR | 18 |
| unknown | 18245 | 3'UTR | 29801 | 3'UTR | 18 |

... (Truncated, full table available in S1)
32 (out of 982) novel junctions that pass SICILIAN from all infection experiments are shown, ordered by number of reads supporting the junction. The 5' and 3' positions of the junctions are indicated, along with their classification as Canonical, L-novel, Internal, or 3' UTR (see main text and methods).

**Table S2.**
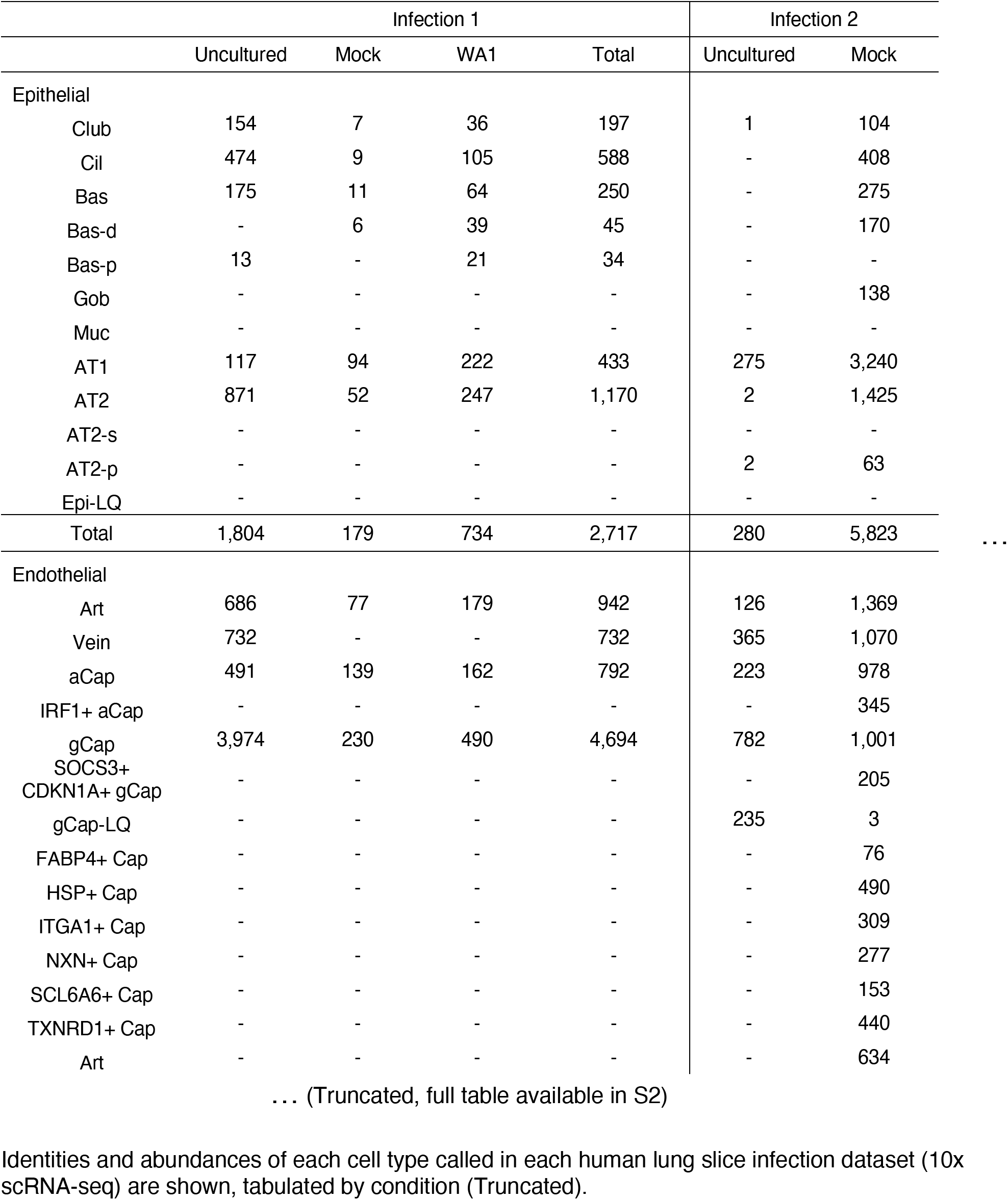
Human lung cell cluster identities and their abundances in each dataset.

|  | Infection 1 |  |  |  | Infection 2 |  |  |
| --- | --- | --- | --- | --- | --- | --- | --- |
|  | Uncultured | Mock | WA1 | Total | Uncultured | Mock |  |
| Epithelial |  |  |  |  |  |  |  |
| Club | 154 | 7 | 36 | 197 | 1 | 104 |  |
| Cil | 474 | 9 | 105 | 588 | - | 408 |  |
| Bas | 175 | 11 | 64 | 250 | - | 275 |  |
| Bas-d | - | 6 | 39 | 45 | - | 170 |  |
| Bas-p | 13 | - | 21 | 34 | - | - |  |
| Gob | - | - | - | - | - | 138 |  |
| Muc | - | - | - | - | - | - |  |
| AT1 | 117 | 94 | 222 | 433 | 275 | 3,240 |  |
| AT2 | 871 | 52 | 247 | 1,170 | 2 | 1,425 |  |
| AT2-s | - | - | - | - | - | - |  |
| AT2-p | - | - | - | - | 2 | 63 |  |
| Epi-LQ | - | - | - | - | - | - |  |
| Total | 1,804 | 179 | 734 | 2,717 | 280 | 5,823 | ... |
| Endothelial |  |  |  |  |  |  |  |
| Art | 686 | 77 | 179 | 942 | 126 | 1,369 |  |
| Vein | 732 | - | - | 732 | 365 | 1,070 |  |
| aCap | 491 | 139 | 162 | 792 | 223 | 978 |  |
| IRF1+ aCap | - | - | - | - | - | 345 |  |
| gCap | 3,974 | 230 | 490 | 4,694 | 782 | 1,001 |  |
| SOCS3+<br>CDKN1A+ gCap | - | - | - | - | - | 205 |  |
| gCap-LQ | - | - | - | - | 235 | 3 |  |
| FABP4+ Cap | - | - | - | - | - | 76 |  |
| HSP+ Cap | - | - | - | - | - | 490 |  |
| ITGA1+ Cap | - | - | - | - | - | 309 |  |
| NXN+ Cap | - | - | - | - | - | 277 |  |
| SCL6A6+ Cap | - | - | - | - | - | 153 |  |
| TXNRD1+ Cap | - | - | - | - | - | 440 |  |
| Art | - | - | - | - | - | 634 |  |

... (Truncated, full table available in S2)
Identities and abundances of each cell type called in each human lung slice infection dataset (10x scRNA-seq) are shown, tabulated by condition (Truncated).

**Table S3.**
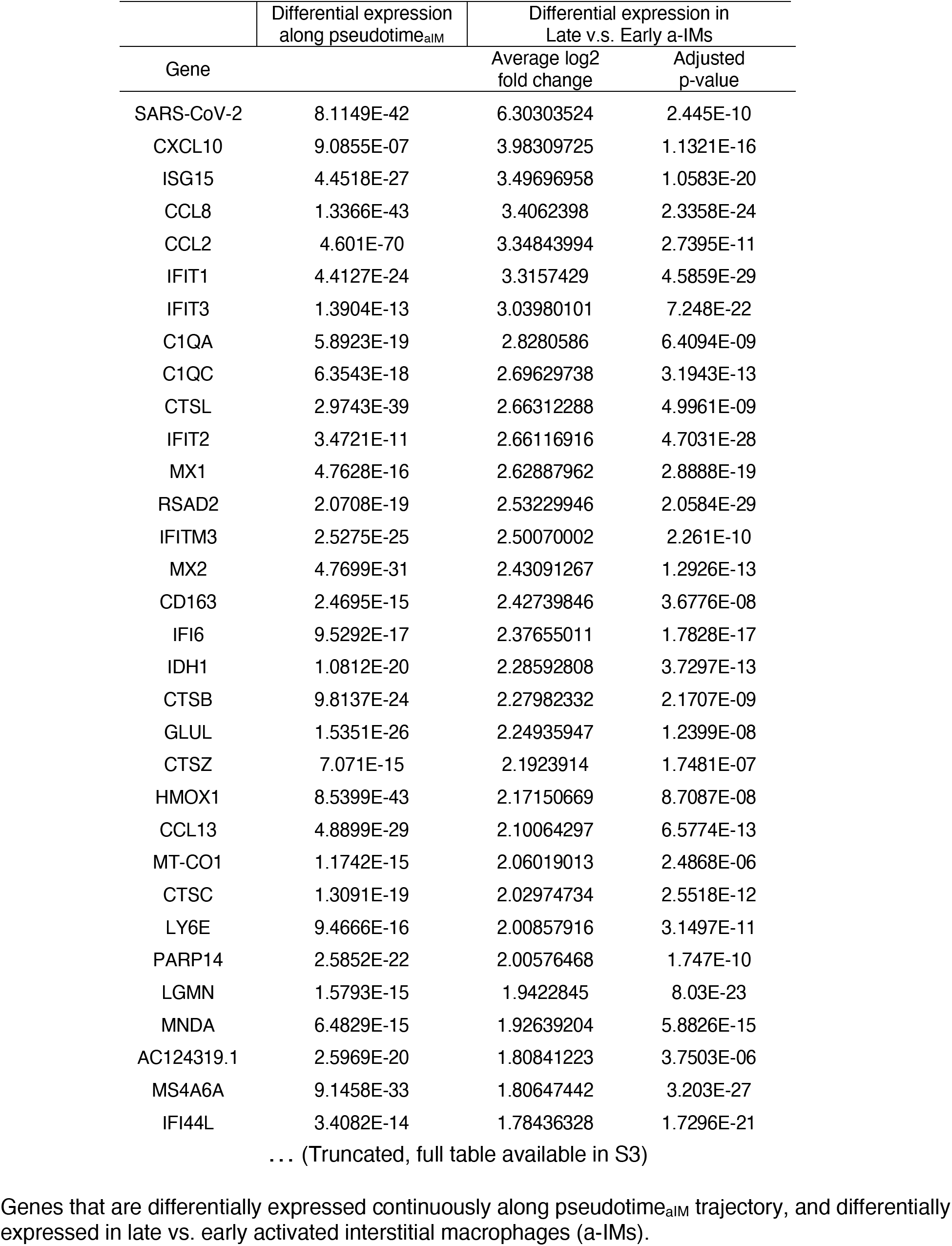
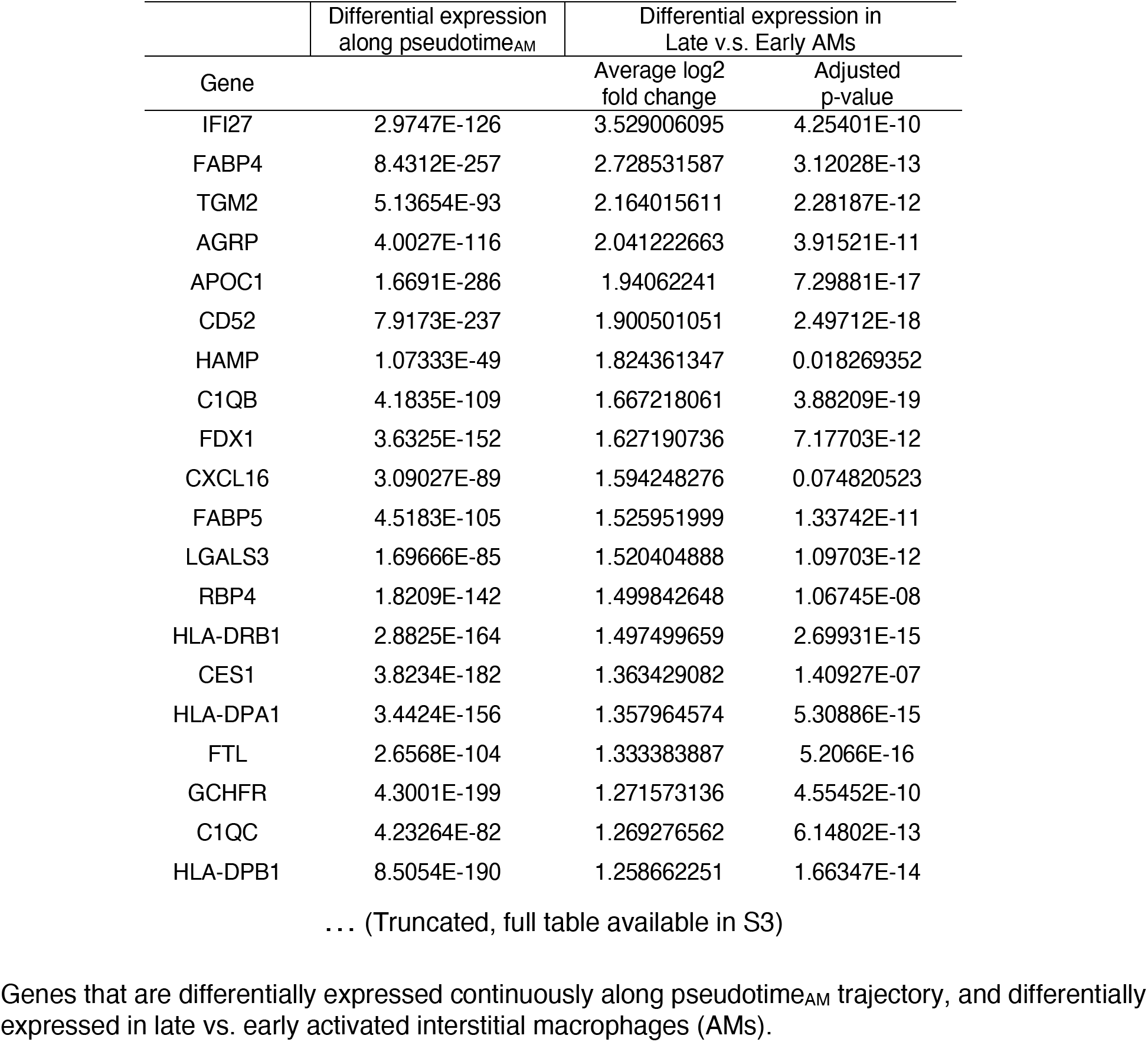
Differentially expressed genes along infection pseudotime trajectory for alveolar and interstitial macrophages.

**Table S4. Clinical summaries of donors or patients of surgical resection**

